# The RNA helicase DDX53 (CAGE) contributes to RNA metabolism in a human germ cell model

**DOI:** 10.64898/2026.02.03.703580

**Authors:** Agata J. Barszcz, Katarzyna Tutak, Agnieszka Malcher, Małgorzata Dąbrowska, Joanna Zyprych-Walczak, Erik Dassi, Erkut Ilaslan, Marta Olszewska, Dominik Cysewski, Michał Hrab, Tomasz Kolanowski, Alexander Yatsenko, Maciej Kurpisz, Natalia Rozwadowska

## Abstract

*DDX53* (DEAD-box helicase 53, known also as CAGE) is an intronless gene on the X chromosome, which expression shows strong testis specificity. It belongs to the group of cancer-testis (CT) antigens, with most studies to date focusing on its role in cancer, but the precise biological function of DDX53 remains unclear. Previous reports identifying rare *DDX53* variants in infertile men provided the rationale for investigating the role of DDX53 in the context of human spermatogenesis. By using the human seminoma cell line (TCam-2) as an *in vitro* male germline model, we aimed to investigate the function and molecular targets of DDX53 protein. Our eCLIP and RNA-seq data show that DDX53 protein directly interacts with numerous RNA molecules, drives transcriptome changes in human cells, and is involved in alternative splicing of RNA. Moreover, we identified potential DDX53 protein interactors using Co-IP-MS approach. Subcellular localization analysis by confocal microscopy indicated a predominantly cytoplasmic distribution of DDX53, with partial nuclear presence in TCam-2 cells. We also identified DDX53-positive structures that may correspond to germ granule-like assemblies, although their precise nature remains to be determined. Additionally, we confirmed DDX53 presence in human testis using a specific, commercially available anti-DDX53 antibody. Our data indicate that DDX53 protein acts as a regulator of RNA metabolism in human cells. Collectively, we show that DDX53 participates in transcriptome regulation (including splicing) in male germ cells and exhibits transcriptome-wide RNA interactions, but its wider biological role remains to be clarified.

## 1. Introduction

The *DDX53* (DEAD-box helicase 53; known also as CAGE (cancer-associated gene protein, or cancer/testis antigen 26)) is an intronless gene, located in the Xp22.11 region, and rare variants within this gene have been recently described as possibly associated with spermatogenesis failure based on whole genome/exome sequencing (WGS/WES) genetic screening of infertile patients [1,2]. Published phylogenetic trees of DDX53 demonstrated a high degree of sequence homology with most primate species [3,4]. Interestingly, although it displays an evolutionary relationship with rodent species (such as the European rabbit or guinea pig), it is not expressed in mice. Considering that species-specific differences in the regulation and function of *DDX53* may exist, a comprehensive characterization of this protein and the construction of a suitable research model remain challenging.

Importantly, the DDX53 protein belongs to the group of cancer-testis antigens (CTAs), defined as proteins that appear to be present only in germ cells and trophoblasts, but aberrantly expressed in various cancer types [5,6]. Currently, CTAs are considered as potential therapeutic targets or novel biomarkers. Although DDX53 expression seems to be restricted to the testis and cancers [7–9], previous reports indicate its expression also in the brain and normal colonic mucosa [4,10]. Studies that investigated the role of DDX53 in the context of cancer reported that it is associated with well known cancer hallmarks [11] such as angiogenesis [12,13], proliferation [14–16], invasion [12,15,17] and metastasis [18–20]. It has been shown that *DDX53* expression is regulated by hypomethylation of its promoter [21] and microRNAs (such as miR-200b, miR-22, and miR-30a) [12,22,23]. Moreover, *DDX53* possesses oncogenic potential and it has been suggested that hypomethylation of *DDX53* may be associated with tumorigenesis progression and may therefore serve as diagnostic marker in cancer [14,21]. DDX53 has also attracted attention because its presence confers resistance to anti-cancer drugs [12,15–17,19,20,23–27]. A note of caution is warranted when interpreting existing data on DDX53. Our observations suggest that some antibodies commonly used to detect DDX53 may show partial cross-reactivity with other DDX family members, including DDX43 (unpublished data). In addition, certain antibodies previously applied have since been withdrawn, raising further questions about their specificity. These factors highlight the need for careful evaluation of antibody-based results and cautious interpretation of conclusions drawn from earlier studies.

According to the Human Protein Atlas (HPA) [28] *DDX53* is expressed predominantly in the human testis tissue, specifically, its presence on mRNA level has been detected in spermatogonia, spermatocytes, and early spermatids where it shows the highest expression.

The DDX53 contains a DEAD [Asp-Glu-Ala-Asp] box domain which is known to contribute to the binding site of RNA molecules as well as ATP hydrolysis and has been resolved by X-ray crystallography [29]. This protein is thought to have an RNA helicase activity and proteins belonging to this family are known to have essential roles in RNA metabolism (including splicing) and have emerged as crucial players in various stages of germ cell development and not only [30–33]. Notably, the dynamic participation of DEAD-box RNA helicases in RNA metabolism shapes the transcriptomic landscape and thus may influence cell fate [34]. Members of the DEAD-box helicases, for example DDX3Y, DDX25, and DDX4 (VASA), have well-documented and significant roles in germline generation, spermatogenesis, and male infertility [30].

Although great interest has been placed on CTAs in cancer research, their functions in germ cells of the testis remain poorly understood [35,36]. To our knowledge, this is the first report on the new candidate gene for non-obstructive azoospermia (NOA) [2], the *DDX53*, investigating its potential functions as well as protein and RNA interactors in the context of spermatogenesis. In this study, we examined the role of DDX53 using the human seminoma cell line (TCam-2) which shares a similar gene expression profile with human primordial germ cells (hPGCs, the precursor cells of the germline) [37] and could serve as an *in vitro* male germline model. Our data provide an overview of DDX53-associated RNA interactions in a human germ cell model, contributing to the current understanding of its potential involvement in male gamete differentiation and offering a resource for future studies in cancer-related contexts.

## 2. Materials and Methods

### 2.1. Cell line

All experiments were performed using the human testicular cancer cell line (TCam-2) which was gifted from Prof. Sohei Kitazawa (Division of Molecular Pathology, Kobe University, Japan) [38,39]. The TCam-2 cells were cultured in Roswell Park Memorial Institute 1640 Medium (RPMI 1640 Medium, GlutaMAX™ Supplement Thermo Fisher Scientific, 61870044) supplemented with 10% heat-inactivated Fetal Bovine Serum (Cytiva, HYCLSV30160.03) and 1% Antibiotic-Antimycotic solution (100X) (Thermo Fisher Scientific, 15240062) at 37 °C and in a humidified 5% CO_2_ incubator.

### 2.2. TCam-2 cell line modification by lentiviral transduction

The TCam-2 cells were modified using the lentiviral transduction (3rd generation packaging system) according to the procedure described previously [40]. To generate lentiviruses, the following plasmids were used: pRSV-Rev (Addgene, #12253), pMDLg/pRRE (Addgene, #12251), pMSCV-VSV-G (Addgene, #14888), and pLV[Exp]-Puro-EF1A>EGFP/3xFLAG (Vector Builder, Vector ID VB180409-1201grs) or pLV[Exp]-Puro-EF1A>hDDX53[NM_182699.3]/3xFLAG (Vector Builder, Vector ID VB180321-1148yrw) in ratio 1:1:1:2. After 72h, post transfection pseudoviral particles were collected and filtered through 0.45 μm (Millipore, SLHVR13SL). Viral aliquots were frozen and stored at −80 °C. Next 1 × 10^5^ TCam-2 cells were incubated with the viral supernatant in the presence of polybrene (final concentration 4μg/ml, Sigma-Aldrich, TR-1003-G) for 48h. After incubation with the pseudoviral particles, selection with puromycin (final concentration 1 ug/ml, Thermo Fisher Scientific, A1113803) was initiated and continued for 4 days in order to obtain cells stably expressing the DDX53-FLAG and GFP-FLAG (as control).

### 2.3. Transient transfection

The TCam-2 cells were transfected with plasmids listed below using jetPRIME transfection reagent (Polyplus, 114-15). For immunofluorescence experiments, cells were seeded on 15 mm round glass coverslips in 12-well plates and transfected using a mixture prepared according to the manufacturer’s protocol (0.8 μg plasmid, 1.6 μl reagent, 75 μl buffer; 75 μl per well). For other experiments, cells were cultured in 150 mm dishes and transfected with a mixture prepared according to the same protocol (31.5 μg plasmid, 63 μl reagent, 1.3 ml buffer; 1.3 ml per dish) [40]. Next, the TCam-2 cells were cultured in a complete medium with transfection mixtures for 2 days and then subjected to downstream experiments. Plasmids used: (pLV[Exp]-Puro-EF1A>hDDX53[NM_182699.3]/3xFLAG (Vector Builder, Vector ID VB180321-1148yrw), pLV[Exp]-Puro-EF1A>EGFP/3xFLAG (Vector Builder, Vector ID VB180409-1201grs).

### 2.4. RT-qPCR and statistical analysis

Total RNA was extracted using RNeasy Plus Mini Kit (Qiagen, 74136) and DNA residues were removed by TURBO DNA-free™ kit (Thermo Fisher Scientific, AM1907) treatment according to the manufacturer’s protocol. Reverse transcription was conducted with SuperScript IV Reverse Transcriptase kit (Thermo Fisher Scientific, 18090050) with the use of random primers (Promega, C1181) using 1 µg of total RNA for each reaction. Ten ng of cDNA was used for qPCR, performed with PowerUp SYBR Green Master Mix (Thermo Fisher Scientific, A25778) on CFX96 Touch qPCR System (Bio-Rad). All samples were normalized to ACTB and GAPDH according to the geNorm algorithm. Statistical analyses were performed in GraphPad Prism (v8.0.1) using ordinary one-way ANOVA non-parametric test with Tukey’s multiple comparisons test. All primer sequences are listed in Supplementary Table S14.

### 2.5. RNA sequencing and bioinformatic analysis

The human TCam-2 cells transduced with *DDX53-FLAG*, *GFP-FLAG* and non-transduced WT cells were subjected to RNA sequencing (RNA-seq) experiment. The GFP-expressing and WT cells were used as negative controls. Total RNA was isolated and DNase-treated according to the manufacturer’s instructions as it was described above. The cDNA libraries were prepared using TruSeq Stranded mRNA LT kit (Illumina) and then the next-generation sequencing was performed on the Illumina NovaSeq6000 platform, 150 bp paired-end reads with at least 30 M reads per sample. Library preparation and sequencing were performed by Macrogen NGS Service (Korea). All RNA-seq experiments were done in duplicate.

RNA-seq data pre-processing was performed by removing low-quality reads (Q<30) and sequencing adapters using Trim Galore (v0.6.10) (https://github.com/FelixKrueger/TrimGalore). Details of each specific RNA-seq analysis are provided in the following sections.

#### 2.5.1. Differential expression analysis of RNA-seq data

The RNA-seq data were analyzed as we described previously [40]. The pre-processed RNA-seq reads were mapped to the human reference genome (GRCh38/hg38, Gencode v44) using STAR (v2.7.11a) [41] (--outSAMtype BAM SortedByCoordinate --quantMode GeneCounts). BAM files were sorted and indexed using samtools (v1.19.2) [42]. Read counts were then normalized by DESeq2 R package [43]. Lowly expressed genes (with less than 0.1 mean CPM in each condition i.e. DDX53, GFP or WT) were excluded from further analysis. The differential expression analysis was then performed to compare the ‘DDX53 vs. GFP’ and the ‘DDX53 vs. WT’ conditions using DESeq2. Significant differentially expressed genes (DEGs) were selected based on adjusted p-value (FDR) < 0.05 and |log2FC| ≥ 1. As the final list of genes we intersected the significant DEGs of the two comparisons (‘DDX53 vs. GFP’, ‘DDX53 vs. WT’), and we selected only those genes with the same trend (upregulation or downregulation) in both comparisons.

The Gene ontology (GO) and pathway enrichment analyses were performed with the topGO and clusterProfiler [44] R packages, using a 0.05 threshold on Benjamini-Hochberg (BH) adjusted p-value.

#### 2.5.2. Differential splicing analysis of RNA-seq data

We employed the Salmon tool (v1.10.2) [45] to quantify transcript-level expression (TPM) in each condition (DDX53, GFP, and WT) and aligned the pre-processed RNA-seq reads to the reference transcriptome (Gencode v44). Next, we used the SUPPA2 tool (v2.3) [46,47] to perform a differential splicing analysis, setting a threshold of p≤0.05 and |ΔPSI|≥0.15. To identify potentially more specific splicing events, the final list of differentially spliced genes (DSGs) was prepared by doing an intersection between the significant predicted splicing events from the two comparisons (‘DDX53 vs. GFP’ and ‘DDX53 vs. WT’). During the SUPPA analysis, the WT and GFP samples were set always as Cond1 and DDX53 as Cond2. The Gene ontology and pathway enrichment analyses were performed with the topGO and clusterProfiler [44] R packages, using a 0.05 threshold on BH adjusted p-value.

### 2.6. DDX53 eCLIP and bioinformatic analysis

Two days after transfection (using the pLV[Exp]-Puro-EF1A>hDDX53[NM_182699.3]/3xFLAG plasmid), TCam-2 cells on multiple 150 mm cell culture dishes were crosslinked at 254-nm UV with an energy setting of 400 mJoules/cm2 (Analytik Jena, CL-3000). Two biological replicates of TCam-2 cells expressing the DDX53-FLAG were used in eCLIP-seq performed by Eclipse BioInnovations (USA). Briefly, the cells were lysed, RNase I treated, and protein-RNA complexes were immunoprecipitated with anti-FLAG antibody coupled to magnetic beads. Sequencing was performed as SE72 on the NextSeq platform (Illumina). The eCLIP experiment was performed according to Eclipse BioInnovations’ standard RBP-eCLIP pipeline as previously described [48]. Raw sequencing reads were trimmed of UMIs and adapters, filtered out of reads that mapped to repeat elements, aligned to the reference genome (hg38) with STAR (v2.7.11a), and filtered of PCR duplicates. Final peaks for each sample were filtered for enrichment over the input (|log2FC| ≥ 3) and statistical significance (adjusted p-value ≤ 0.001). De novo motif analysis for DDX53 was performed with HOMER (v4.11).

### 2.7. Co-immunoprecipitation and mass spectrometry (Co-IP-MS)

3 × 10^6^ of TCam-2 cells after transfection with plasmids coding for DDX53-FLAG or GFP-FLAG (used as a negative control) were crosslinked for 10 mins at RT using 0.3% formaldehyde (Thermo Fisher Scientific, 28906), then quenched with 125 mM glycine (BioShop, GLN001) in PBS for 5 min at RT and 5 mins on ice, and finally lysed with 1ml of RIPA buffer (Sigma-Aldrich, R0278) supplemented with protease inhibitors (Thermo Fisher Scientific, 87786) at 4°C for 30 min on rotator. Total protein lysates were centrifuged (4°C, 10 min, 11 000 x g) and 1 ml was incubated with 40µl of anti-FLAG M2 magnetic beads (Sigma-Aldrich, M8823) at 4°C for 3 h with gentle rotation. After IP reaction, beads were washed three times with ice-cold TBS washing buffer (50 mM Tris HCl, 150 mM NaCl, pH 7.4) and one time with DPBS (Thermo Fisher Scientific, 14190169). Three technical replicates of IP reactions were prepared. For mass spectrometry (MS), beads were snap frozen with liquid nitrogen and processed as previously described [40]. MS analysis was carried out at the Mass Spectrometry Laboratory, IBB PAS. Dried beads were suspended in 50μl of 100mM ABC. Proteins were reduced and alkylated with TCEP/MMTS, followed by overnight digestion with trypsin (Promega, V5111) as described previously [49]. Peptide digest was purified with HLB 96-well plates (Waters, WAT058951), vacuum-dried, dissolved in 98% water (J.T.Baker 4218-03)/ 2% MeCN (J.T. Baker HPLC JT9012)/ 0.1%TFA (Supelco, 80457), and analyzed by an online LC-MS/MS (liquid chromatography coupled to tandem mass spectrometry) using a Nano-Acquity (Waters) UPLC system and QExative Orbitrap mass spectrometer (Thermo Fisher Scientific). Peptides were first loaded onto a precolumn (nanoACQUITY UPLC Trapping Column Waters) using 0.1% formic acid in water as a mobile phase and then transferred to a nano-column (nanoACQUITY UPLC BEH C18 Column (75 μm inner diameter; 250 mm long; Waters, 186007482)) using an acetonitrile gradient (5–35% AcN in 180 min) in the presence of 0.1% formic acid with the flow rate of 250 nl/min. Each analysis was preceded by three washing runs to ensure there was no cross-contamination from previous samples. The column outlet was directly coupled to the ion source of the spectrometer working in the regime of data-dependent acquisition MS to MS/MS switch. Peptides were directly eluted into the ion source of the mass spectrometer. MS data were acquired in the m/z range of 300–2000. The resulting spectra were searched against the human UniProt reference proteome (79,052 entries) using MaxQuant (v1.5.3.0) [50]. The following parameters were applied: a match between runs (match time window, 0,7 min; alignment time, 20 min); enzyme, trypsin/p specific; max missed, 2; minimal peptide length, 7 aa; variable modification, methionine, and proline oxidation; fixed, cysteine alkylation (Methylthio); main search peptide tolerance, 4.5 ppm; protein FDR 1%. Finally, 36 proteins identified in each DDX53 replicate but not present in negative controls (GFP) were classified as potential DDX53 protein interactors. Gene ontology analysis among identified proteins was performed in Cytoscape (v3.9.1) using ClueGO plugin (v2.5.9), 4% enrichment cutoff was used meaning that the number of genes should represent at least 4% of total genes in a given process. STRING analysis of protein-protein interaction (PPI) networks was conducted using STRING website (v12.0), a 0.4 cutoff score was used which means interactions below this value were filtered out.

### 2.8. Immunofluorescence of TCam-2 cells

The TCam-2 cells were immunostained according to the procedure described previously [40]. Briefly, cells were washed with DPBS (Thermo Fisher Scientific, 14190169), fixed with 4% paraformaldehyde (Boster Bio, AR1068) at 4 °C for 15 min, washed again with DPBS, permeabilized with 0.55% Triton X-100 in DPBS for 30 mins at RT and blocked for 1h at RT using blocking solution (10% of suitable serum derived from the host animal species of the secondary antibody, diluted in 0.55% Triton-X100 in DPBS). After overnight incubation at 4 °C with primary antibodies diluted in the blocking solution and subsequent washing with DPBS, the cells were incubated with secondary antibodies diluted in 0.55% Triton X-100 in DPBS for 1 h at RT in the dark. After washing, cells were mounted in Fluoroshield with DAPI mounting medium (Sigma-Aldrich, F6057).

Primary antibodies used: rabbit DDX53 (CAGE) 1:100 (Sigma-Aldrich SAB1402037), mouse anti-FLAG 1:500 (Sigma-Aldrich, F1804), rabbit anti-DYKDDDDK 1:2000 (GenScript, A00170), mouse anti-THOC1 1:1000 (Abcam, ab487), rabbit anti-SCML2 1:500 (Sigma-Aldrich, HPA064713), rabbit anti-ACLY 1:1200 (Thermo Fisher Scientific, PA5-29495). Secondary antibodies used: goat anti-mouse Alexa Fluor 594 1:500 (Thermo Fisher Scientific, A-11005), goat anti-rabbit Alexa Fluor 488 1:1000 (Abcam, ab150077), donkey anti-mouse Alexa Fluor 488 1:1000 (Thermo Fisher Scientific, A-21202), goat anti-rabbit Alexa Fluor 594 1:500 (Thermo Fisher Scientific, A-11012).

### 2.9. Confocal microscopy and image analysis

All images were acquired using Leica DMi8 confocal microscope with Leica STELLARIS STED system equipped with Leica HyD photodetectors (Leica Microsystems). Immunofluorescence images were captured and processed using Leica Application Suite X (LAS X) software (v4.4.0.24861) and the HC PL APO CS2 objective (100X magnification, 1.4 NA).

In order to investigate the localization of the DDX53 protein acquired images were analyzed for assessing the percentage of red channel intensities in each nucleus to the whole cell by manually selecting the nucleus area in each optical section of the height scan (z-stack). Each optical section was 0.2 um thick and for every one the intensity sum in the nucleus and intensity sum in the whole cell was calculated. For each cell, at least 4 optical sections were selected for analysis. Subsequently, the ratio of intensity sum in the nucleus to the whole cell was assessed by adding values obtained for each optical section. Finally, the average ratio for all cells selected for analysis was calculated.

### 2.10. Immunohistochemistry

Immunohistochemistry (IHC) procedure was performed as we described previously [40]. For the IHC staining, commercial human adult testis paraffin sections were used (Zyagen, HP-401). Slides were incubated in 60°C for one hour. Immunohistochemical stainings were performed using specific DDX53 antibody (Sigma-Aldrich SAB1402037). To determine the appropriate antibody dilution, eliminate false positive results, and reduce the background staining, a series of positive control reactions were performed using slides with normal testis cells present in section derived from patient with seminoma. The immunohistochemistry procedure was performed using EnVisionFlex+ visualization system kit (Dako, Agilent Technologies, K8002). Slides were then dewaxed and epitopes were unmasked in PT-Link device (Dako, Agilent Technologies) with the use of EnVisionFlex Target Retrieval Solution Low pH (Dako, Agilent Technologies, K8005). Tissue sections were then incubated for 20 mins at RT with primary antibody. The antigen–antibody complexes were visualized using EnVision FLEX+ Rabbit LINKER (Dako, Agilent Technologies, K8009) for 20 min at RT and localized using 3-3′diaminobenzidine (DAB) as chromogen. Slides were counterstained with hematoxylin, dehydrated in ethanol, incubated in a series of xylenes, and cover-slipped using mounting medium (Dako, Agilent Technologies). Protein expression was assessed under the Leica DMi8 light microscope.

### 2.11. Western blot

TCam-2 cells were lysed on a rotator for 30 min in RIPA buffer (Sigma-Aldrich, R0278) supplemented with protease inhibitors (Thermo Fisher Scientific, 87786). Samples were subjected to three rounds of sonication at 4°C [20 s on, 30 s off] using Bioruptor Pico (Diagenode, Belgium). Then, protein samples were centrifuged at 15000 x g for 10 min at 4°C and supernatants were mixed with 4X Laemmli Sample Buffer (Bio-Rad, 1610747) with 340 mM DTT (Thermo Fisher, R0861), denatured for 10 min at 95°C, separated on 8% SDS polyacrylamide gels and transferred to nitrocellulose membranes (Bio-Rad, 1620097). Membranes were blocked in 5% non-fat milk in TBS buffer with 0.1% Tween-20 for 1 hour, and incubated overnight at 4 °C with primary antibodies diluted in the blocking solution. Washing steps (3×10min. with TBS buffer with 0.1% Tween 20) were followed by incubation with horseradish peroxidase (HRP)-conjugated secondary antibodies for 1h at RT. Blots were washed twice with TBS buffer with 0.1% Tween-20 for 10 min and twice with TBS for 10 min. The chemiluminescent signal was detected using the Clarity Western ECL Substrate (Bio-Rad, 1705061) using ChemiDoc MP Imaging System (Bio-Rad). Antibodies used in this study were as follows: mouse anti-FLAG 1:1000 (Sigma Aldrich, F1804), mouse anti-GAPDH 1:10 000 (Santa Cruz, sc-365062), mouse anti-TUBB 1:5000 (Abcam, ab131205), mouse anti-SNRNP70 1:5000 (Santa Cruz, sc-390899), rabbit anti-H3C1 1:5000 (Abcam, ab18521), HRP-conjugated anti-rabbit 1:5000 (Abcam, ab6721), HRP-conjugated anti-mouse 1:10 000 (Santa Cruz Biotechnology, sc-516102).

### 2.12. Subcellular fractionation

Protein fractionation of TCam-2 cells (lentiviral-modified and WT) was performed according to the previously described method [51]. Briefly, cell pellets were lysed on ice using 1× lysis buffer (10 mM Tris–HCl pH 8, 300 mM sucrose, 10 mM NaCl, 2 mM MgAc₂, 3 mM CaCl₂, 0.1% Nonidet P-40, 0.5 mM DTT), followed by centrifugation at 1,000×g for 5 min at 4°C to obtain the cytoplasmic fraction. The nuclear pellet was subsequently washed once with glycerol buffer (50 mM Tris–HCl pH 8.0, 25% glycerol, 5 mM MgAc₂, 0.1 mM EDTA, 5 mM DTT). To further separate nucleoplasmic and chromatin fractions, nuclei were lysed in urea buffer (20 mM HEPES pH 7.5, 7.5 mM MgCl₂, 0.1 mM EGTA, 0.3 M NaCl, 1 M urea, 1% Nonidet P-40, 1 mM DTT) and centrifuged at 14,000×g for 10 min at 4°C. All buffers were supplemented immediately before use with 20 U/mL RNaseOUT (Thermo Fisher Scientific, 10777019) and 1× Complete™ EDTA-free protease inhibitor cocktail (Thermo Fisher Scientific, 87786).

## 3. Results

### 3.1. Detection of DDX53 protein in the human testis and generation of in vitro germ cell model with DDX53 overexpression

The gene expression profile analysis, based on independent data set [52,53], showed differences in the expression of *DDX53* in infertile men with idiopathic non-obstructive azoospermia (NOA) compared to men with normal spermatogenesis (Figure 1A). In order to confirm the presence of DDX53 in normal adult human testis we performed immunohistochemical staining for DDX53 using tissue section (Figure 1B). This experiment showed that DDX53 is present in spermatocytes but it is the most visible in spermatids. The detection of DDX53 in normal human testis, lower expression level in azoospermic patients, and possible association of *DDX53* with spermatogenesis failure based on the literature [1,2] encouraged us to investigate its function in the context of spermatogenesis. In our study, we used the human seminoma cell line (TCam-2), which in terms of gene expression resembles human primordial germ cells (hPGC), the embryonic precursors of gametes [37]. For the purpose of investigating the DDX53 protein function, we modified WT TCam-2 cells to obtain cells expressing the FLAG-tagged *DDX53* and FLAG-tagged *GFP* (as a control). Successful experimental model generation was confirmed by RT-qPCR and Western Blot (Figure 1C, D). Moreover, co-immunofluorescence staining using anti-FLAG and specific anti-DDX53 antibodies demonstrated complete signal co-localization coming from the *DDX53-FLAG* (Figure 1E).

**Figure 1.**
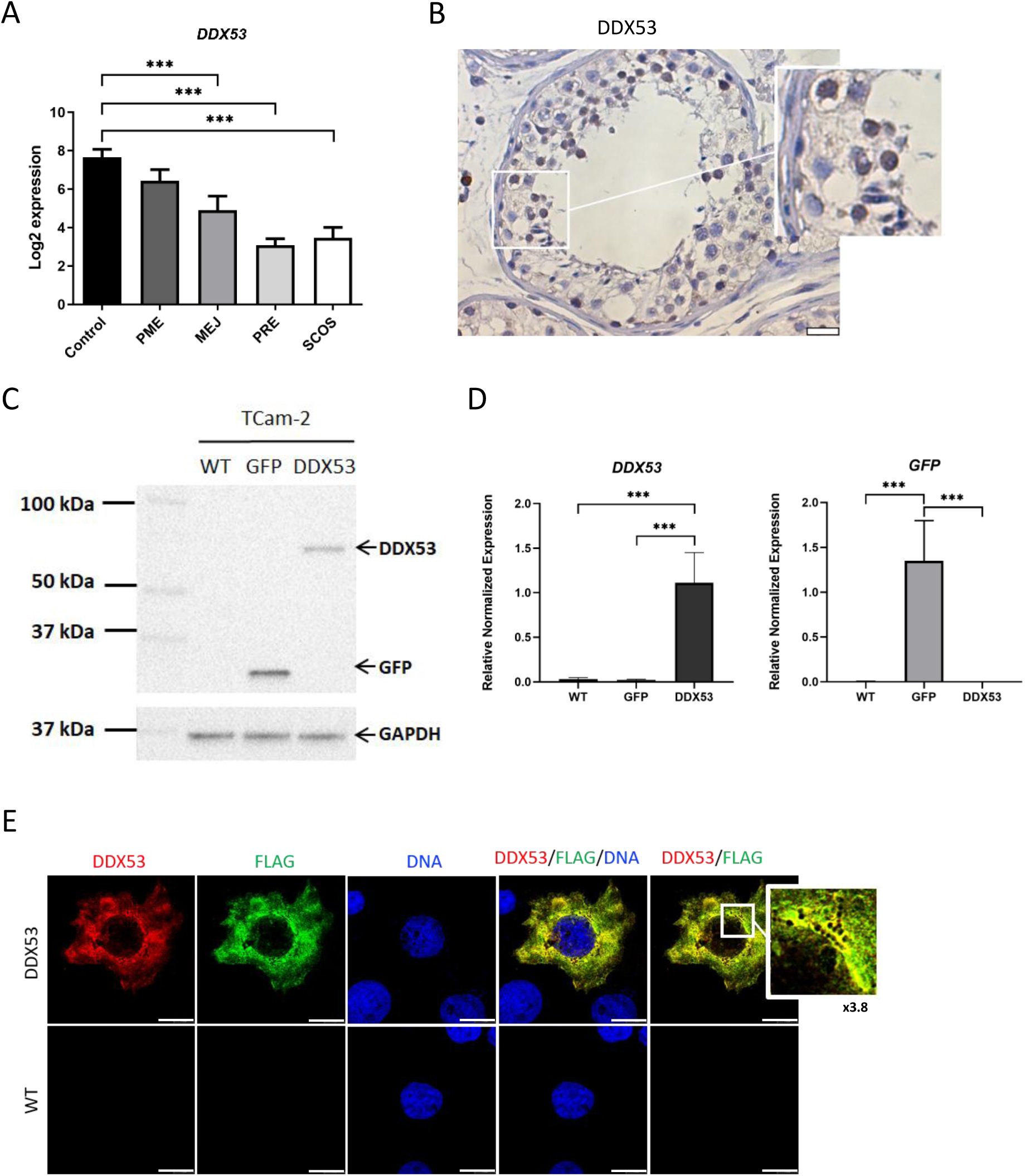
The *DDX53* expression in men with idiopathic non-obstructive azoospermia (NOA), DDX53 in human testis with maintained spermatogenesis, and lentiviral transduction for stable expression of DDX53 in TCam-2 cells. **(A)** Gene expression level of *DDX53* in men with normal spermatogenesis (control, n=5) compared to men with different types of azoospermia (PME - postmeiotic arrest, n = 6; MEJ - meiotic arrest, n = 5; PRE - premeiotic arrest, n = 3; SCOS - Sertoli cell-only syndrome, n = 5) analyzed by microarrays analysis using published data sets [52,53]. Error bars represent standard deviation (SD), ***p < 0.001. The p-values were calculated by ordinary one-way ANOVA non-parametric test with Tukey’s multiple comparisons test. **(B)** Immunohistochemistry of DDX53 using specific DDX53 antibody in the normal adult human testis. Scale bar 25 μm. **(C)** Representative Western Blot analysis for proteins tagged with FLAG (DDX53 and GFP) using TCam-2 cells after transduction and wild type (WT, as control) using FLAG antibody. GAPDH served as a loading control. **(D)** The mRNA expression level of the *DDX53-FLAG* and *GFP-FLAG* in TCam-2 cells modified with lentiviral transduction, and with *GAPDH* and *ACTB* as reference genes. Error bars represent standard deviation (SD) with n = 3, ***p < 0.001. The p-values were calculated by ordinary one-way ANOVA non-parametric test with Tukey’s multiple comparisons test. **(E)** Co-immunofluorescence staining using anti-FLAG and specific anti-DDX53 antibody in the TCam-2 cells expressing DDX53-FLAG and wild type (WT). The zoom window illustrates signal colocalization of DDX53 and FLAG, the magnification factor (×3.8) is indicated below. Scale bar 20 μm.

### 3.2. DDX53 protein regulates the human transcriptome and influences alternative splicing of RNA transcripts

Subsequently, in order to investigate the impact of DDX53 protein overexpression on the TCam-2 transcriptome we performed RNA-sequencing (RNA-seq) using cells stably expressing either the *DDX53-FLAG* or *GFP-FLAG* as well as wild-type (WT) cells (Figure 2A, Supplementary Table S1; Supplementary Figure S1A). Since both the GFP and WT TCam-2 cells were used as controls we focused on genes that were the differentially expressed genes (DEGs) in both comparisons i.e. ‘DDX53 vs. GFP’ and ‘DDX53 vs. WT’ (see Materials and Methods section for more details). Employing a threshold of |log2FC| ≥ 1| and FDR < 0.05, we identified 70 up-regulated genes and 65 down-regulated genes induced by *DDX53* overexpression (Figure 2B, Supplementary Table S2, Supplementary Figure S1B, C). The majority of up-regulated DEGs belonged to protein-coding transcripts, however, in the group of DEGs there were also long non-coding RNAs (lncRNAs) (Figure 2C). In order to validate our RNA-seq results, we selected three genes expressed in human testis (*SERPINF1*, *MZT1*, and *FBN2*) and confirmed their mRNA expression pattern in TCam-2 cells overexpressing *DDX53* via RT-qPCR with respect to both controls, *GFP* and WT cells (Figure 2D, Supplementary Figure S1D).

**Figure 2.**
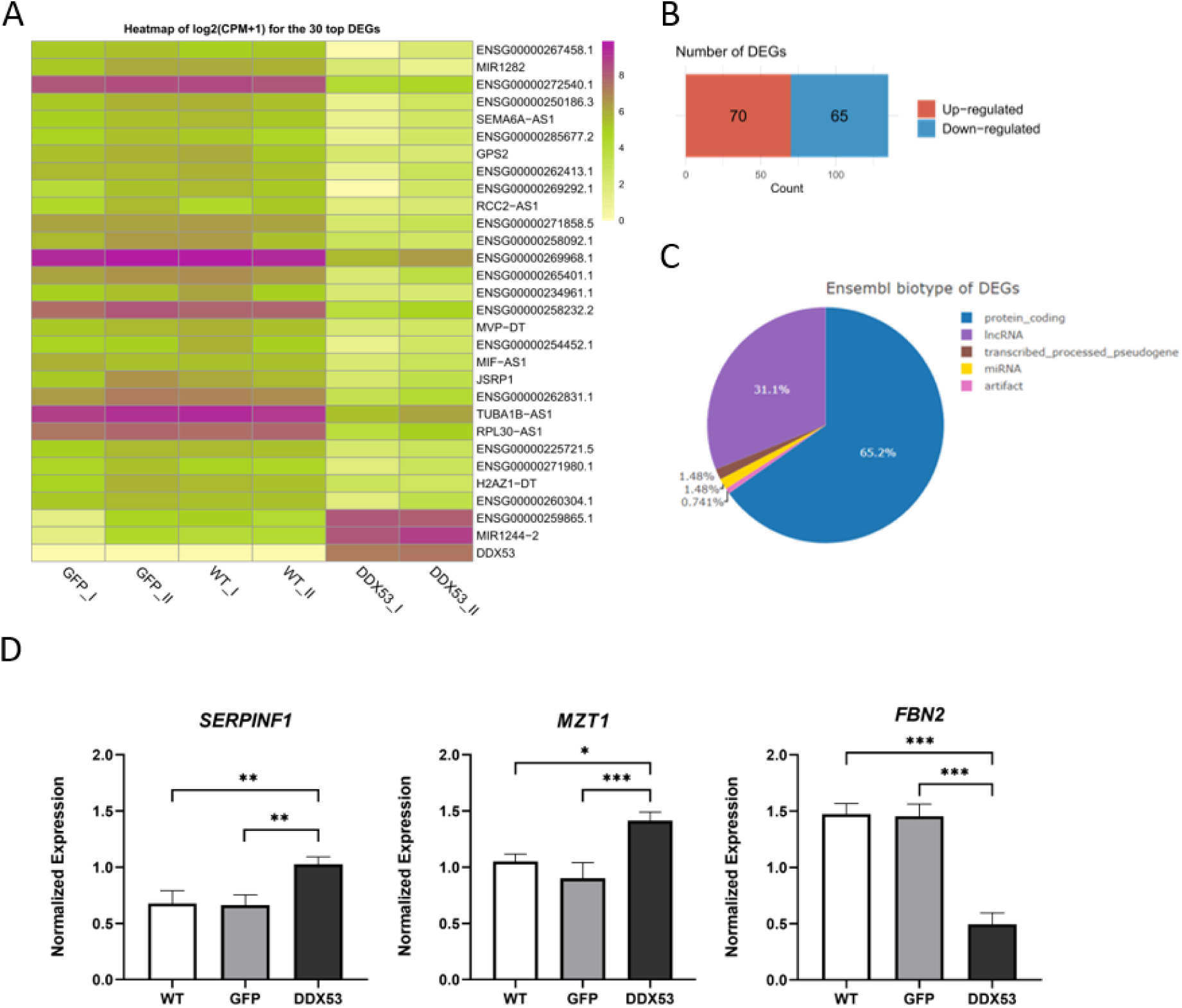
Identification of differentially expressed genes upon *DDX53* overexpression in TCam-2 cells. **(A)** Heatmap of log2(CPM+1) for the top 30 differentially expressed genes (DEGs) based on RNA-seq analysis of TCam-2 cells overexpressing *DDX53-FLAG*, *GFP-FLAG*, and WT (|𝑙𝑜𝑔2𝐹𝐶|≥1, FDR < 0.05). The top 30 DEGs were selected based on the highest |𝑙𝑜𝑔2𝐹𝐶| and the lowest FDR values derived from the ‘DDX53 vs. GFP’ comparison (Supplementary Table S1, 2). **(B)** The number of down-regulated and up-regulated differentially expressed genes (DEGs) based on RNA-seq analysis (Supplementary Table S2), |𝑙𝑜𝑔2𝐹𝐶|≥1, FDR < 0.05. **(C)** Piechart showing the Ensembl biotype of DEGs based on RNA-seq analysis of TCam-2 cells (|𝑙𝑜𝑔2𝐹𝐶|≥1, FDR < 0.05). **(D)** Validation results of selected (*SERPINF1*, *MZT1*, and *FBN2*) DEGs via RT-qPCR confirmed RNA-seq expression pattern from cells overexpressing *DDX53-FLAG*, *GFP-FLAG*, and WT. Error bars represent standard deviation (SD) with n = 3, ***p < 0.001, **p < 0.01, *p < 0.05. The p-values were calculated by ordinary oneway ANOVA non-parametric test with Tukey’s multiple comparisons test.

Due to the fact that some of RNA helicases are thought to participate in RNA splicing, we analyzed the RNA-seq data using SUPPA2 software which allows for the identification of different alternative splicing (AS) events. To detect potentially more specific splicing events, we focused on significant splicing events present in both comparisons (‘DDX53 vs. GFP’ and ‘DDX53 vs. WT’, see Materials and Methods section for more details). According to SUPPA2 analysis, 1270 splicing events in 646 transcripts (p≤0.05 and |ΔPSI|≥0.15) were predicted to occur in TCam-2 cells upon expression of DDX53 protein (Supplementary Table S3, 4). The majority of AS events were alternative first exon (n=666) and exon skipping (n=253) (Figure 3A). Then we extracted the transcripts associated with human male gamete generation by using the genes list annotated to the GO term (GO:0048232) and compared them with differentially spliced genes (DSGs) and DEGs. This integrative analysis identified 24 genes, with *PAFAH1B1*, *BAX*, *CDK16*, *SEPTIN7*, *RAN* and *YTHDF2*, among others, that potentially were differentially spliced due to DDX53 protein presence in TCam-2 cells (Figure 3B, Supplementary Table S5). Gene ontology (GO) enrichment analysis performed among the DSGs (Supplementary Table S6) showed that they are associated with terms such as spindle microtubule, mitotic cell cycle process, chromosome, DNA-templated transcription, sex chromosome but also regulation of RNA splicing, mRNA transport and mRNA processing. To further explore the potential involvement of DDX53 targets in spermatogenesis, the Reactome pathway analysis was carried out among the DSGs. This identified terms such as Nonsense-Mediated Decay (NMD), mRNA splicing, and translation, pathways crucial during the differentiation of germ cells (Supplementary Figure S2; Supplementary Table S6).

**Figure 3.**
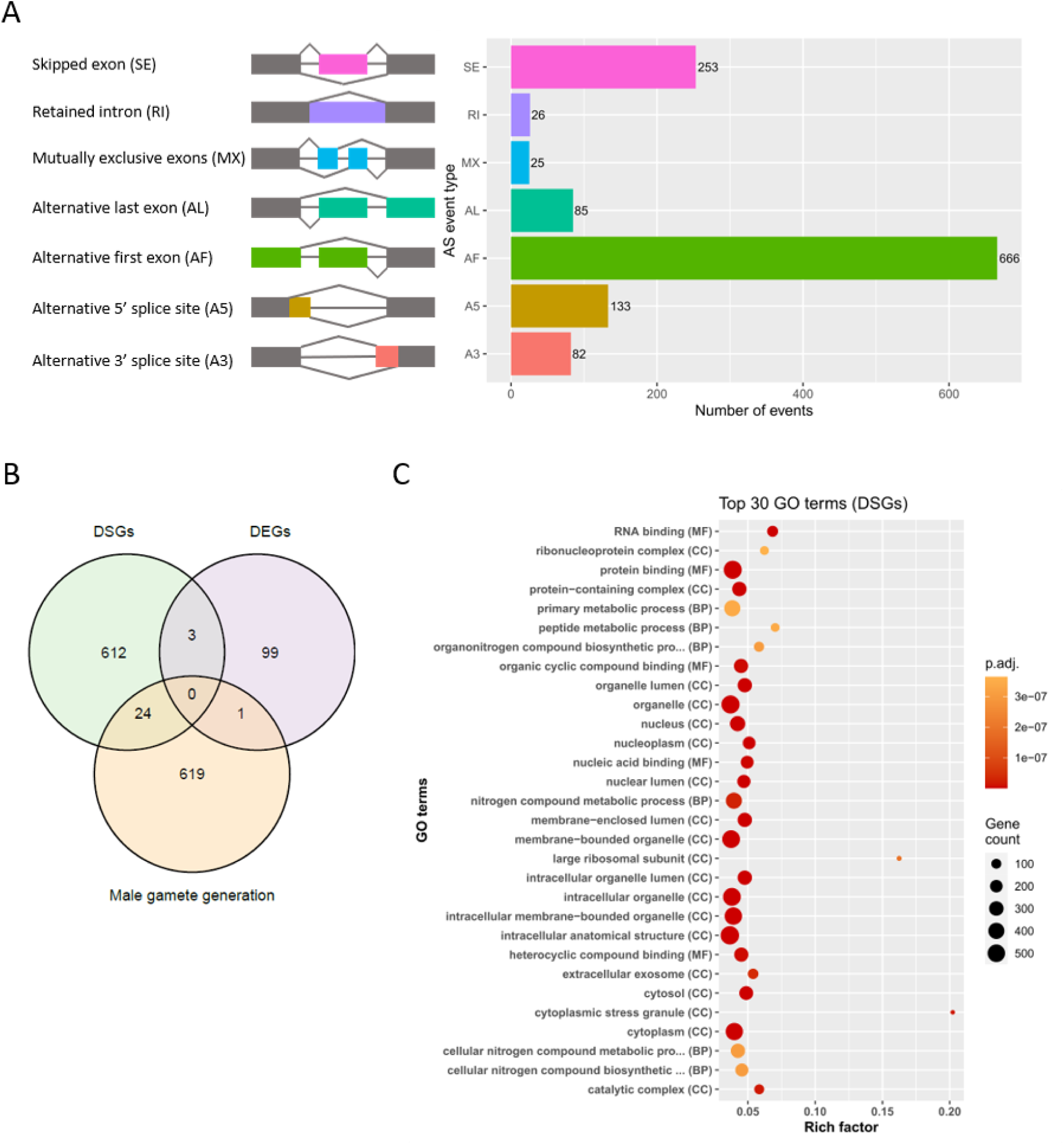
Global identification of alternative splicing (AS) events upon *DDX53* overexpression. **(A)** The total number (1270) of differential alternative splicing (AS, p≤0.05 and |𝑑𝑃𝑆𝐼|≥0.15) events detected by SUPPA2 analysis of RNA-seq data from TCam-2 cells with *DDX53-FLAG* overexpression (Supplementary Table S4). **(B)** Venn diagram illustrating the total number of DSGs and DEGs based on RNA-seq analysis compared to the list of genes associated with human male gamete generation (Supplementary Table S5). The male gamete generation genes list annotated to GO:0048232 was retrieved from AMIGO2 (accession date 2024-02-02). Intersection was prepared using gene symbols. **(C)** GO terms significantly (adjusted p-value < 0.05) enriched for the differentially spliced genes (DSGs) due to overexpression of the DDX53 protein in TCam-2 cells. For the complete list of GO terms see Supplementary Table S6.

### 3.3. DDX53 protein is predominantly cytoplasmic but also exhibits partial nuclear localization in human TCam-2 cells

Previous studies have reported positive DDX53 chromatin immunoprecipitation (ChIP) assay signals, suggesting its binding at promoter regions of protein-coding genes and microRNAs [12,15,23,27,54]. Surprisingly, in our hands, performed ChIP-seq based on FLAG-tagged protein did not show any DNA sequences enrichment (unpublished data). This discrepancy may reflect technical differences between studies, including potential limitations related to the specificity or performance of the antibodies used to detect DDX53 in previous reports. However, an indirect or transient association of DDX53 with chromatin, potentially mediated through protein–protein interactions, cannot be excluded. To examine the localization of DDX53 in human TCam-2 cell line, we performed a series of immunofluorescence stainings using FLAG antibody in cells with transient overexpression of the DDX53 protein. The detailed analysis of DDX53 localization using confocal microscopy and confirmed its presence not only in the cytoplasm but also in the nucleus of TCam-2 cells, albeit at a very limited level (Figure 4A). We calculated the percentage of DDX53 protein presence in the nucleus for each cell separately and obtained the average nuclear localization with respect to the whole cells, which remained low, reaching 11% (Figure 4B). Our immunofluorescence staining images demonstrated similar DDX53 staining pattern consistent with previous report [27]. Additionally, to test the conclusion regarding the partial nuclear localization of DDX53, we performed subcellular fractionation using WT and DDX53-expressing TCam-2 cells (Supplementary Figure S3A). This experiment showed that DDX53 is predominantly localized in the cytoplasm, with a small fraction present in the nucleus of TCam-2 cells. We acknowledge that these findings are based on a single cell line and that it would be valuable to examine DDX53 cellular localization in additional cell lines.

**Figure 4.**
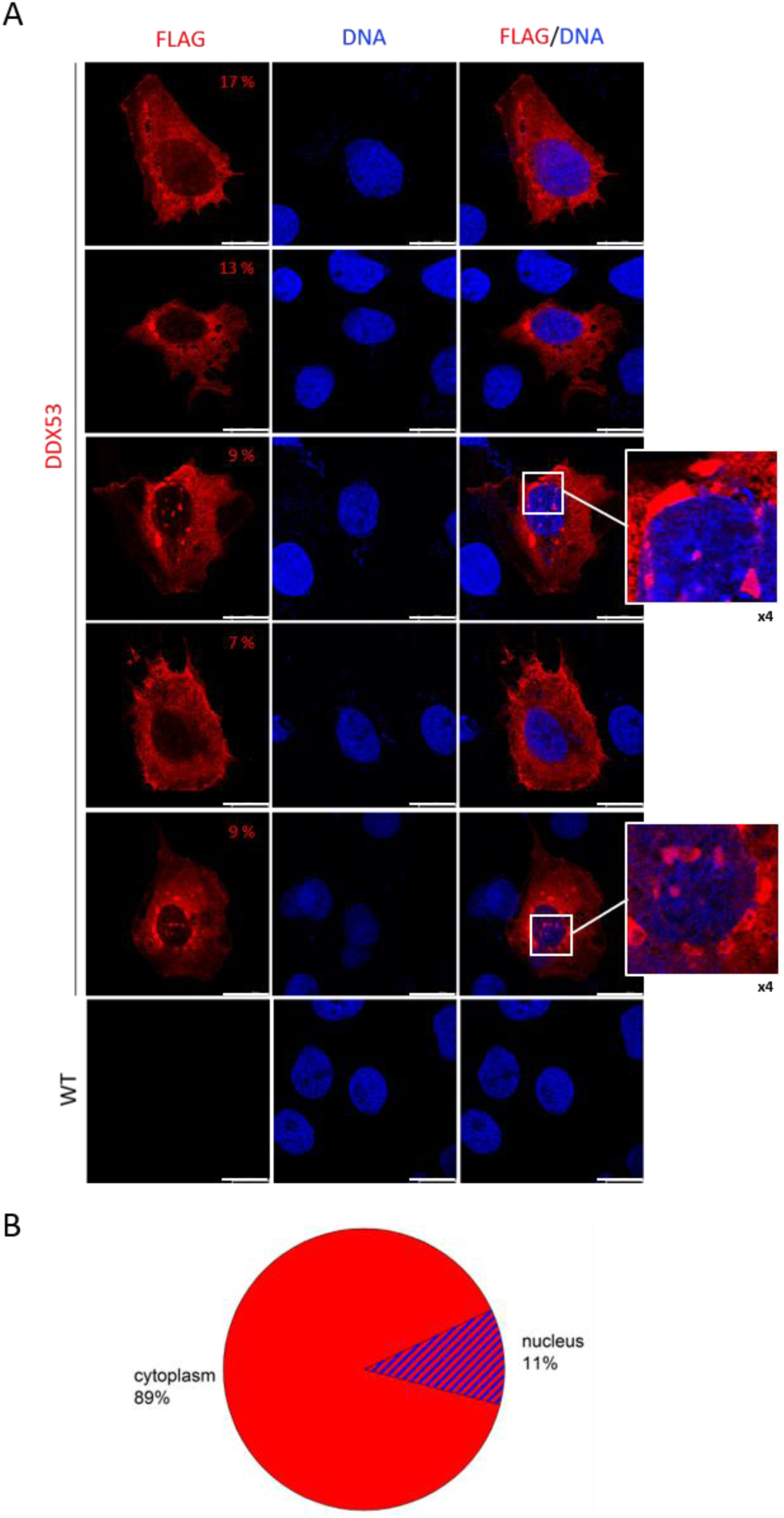
DDX53 protein is predominantly expressed in cytoplasm, but is also present in the nucleus of TCam-2 cells. **(A)** Confocal immunofluorescence staining images of FLAG used for the analysis of DDX53 localization in TCam-2 cells after transfection with a plasmid coding for *DDX53-FLAG*. The numbers in the corner represent the percentage of red channel intensity in the nucleus with respect to the whole cell (for each cell separately). Magnified insets show DDX53-positive condensates; the magnification factor (×4) is indicated below. Immunostaining for FLAG on TCam-2 WT cells was performed as a negative control. Scale bar 20 μm. **(B)** Piechart demonstrating DDX53 localization in a human testicular cancer cell line (TCam-2). The percentage was calculated as the average (from 5 cells shown in panel **A**) of red channel intensities in each nucleus with respect to the whole cell.

Interestingly, using immunofluorescence staining and confocal imaging, we detected DDX53-positive structures that may resemble “germ granules”, nonmembrane-bound organelles unique to the germline. A visible foci of DDX53-positive signals were present only in a subset of TCam-2 cells (Figure 4A and Figure 7 B,C), typically had perinuclear localization, but in certain instances appeared to localize also within the nucleus (intranuclear granules). Germ granules are evolutionarily conserved structures composed of RNAs and RNA-binding proteins that participate in posttranscriptional regulation in developing male germ cells and have essential role in the control of spermatogenesis [55–57]. Although this is an interesting observation, a more in-depth characterization would be essential to determine this with certainty.

### 3.4. Transcriptome-wide discovery of RNAs directly interacting with DDX53 by eCLIP

As DEAD-box domain, present in DDX53 protein, is known to contribute to the binding of RNA molecules, we performed the enhanced crosslinking and immunoprecipitation followed by sequencing (eCLIP-seq). For this purpose, TCam-2 WT cells were transfected with vector encoding *DDX53-FLAG* and expression of DDX53 was confirmed on both RNA and protein levels (Supplementary Figure S3B,C). The eCLIP experiment was performed using two biological replicates of TCam-2 cells transfected with *DDX53-FLAG* (Figure 5A) and eCLIP data were analyzed using very conservative cutoffs, |log2FC| ≥ 3 and adjusted p-value ≤ 0.001. This approach allowed for the identification of ∼1,000 DDX53-binding peaks in over 500 human RNA transcripts, detected in each sample (Supplementary Table S7). Further analysis of DDX53-binding sites (Figure 5B-D) focused on 134 transcripts with reproducible peaks identified across DDX53 replicates. However, as protein–RNA interactions are dynamic, DDX53-binding sites on specific RNAs may differ between replicates (e.g. LIN28B, Supplementary Table S7). Therefore, we prepared the extended list of 181 DDX53-bound transcripts detected in both replicates, regardless of the exact binding site position (Supplementary Table S8). The majority of enriched peaks were found on CDS, 3’UTRs, and distal introns (Figure 5B-D). Additionally, HOMER results showed enrichment of CA motifs (Figure 6A). In order to identify transcripts of differentially spliced genes (DSGs) that were directly bound by DDX53 further analysis was conducted (Figure 6B, Supplementary Table S9). This led to identification of twelve transcripts that might be direct targets of DDX53 protein, *REST*, *TFAM*, *NCBP2*, *CAMSAP2*, *LRP1*, *LRRC8A*, *THAP12*, *LRIG3*, *ESCO1*, *CDH2*, *PEG3*, and *ZNF480*. Subsequently, performed gene ontology (GO) analysis of the detected RNAs revealed that the DDX53 targets are involved in processes such as regulation of RNA metabolic process, transcription by RNA polymerase II, DNA-templated transcription, DNA-binding transcription factor activity, but also cell-cell junction, cell differentiation, and cell population proliferation (Figure 6C, for the complete list of GO terms see Supplementary Table S10). Moreover, evidence for DDX53 binding to its own mRNA was observed, aligning with self-regulatory mechanisms reported for other RNA-binding proteins [58–60]. In conclusion, eCLIP-seq experiment confirmed the functional identity of DDX53 as an RNA-binding protein and identified its RNA targets, highlighting its impact on human transcriptome regulation.

**Figure 5.**
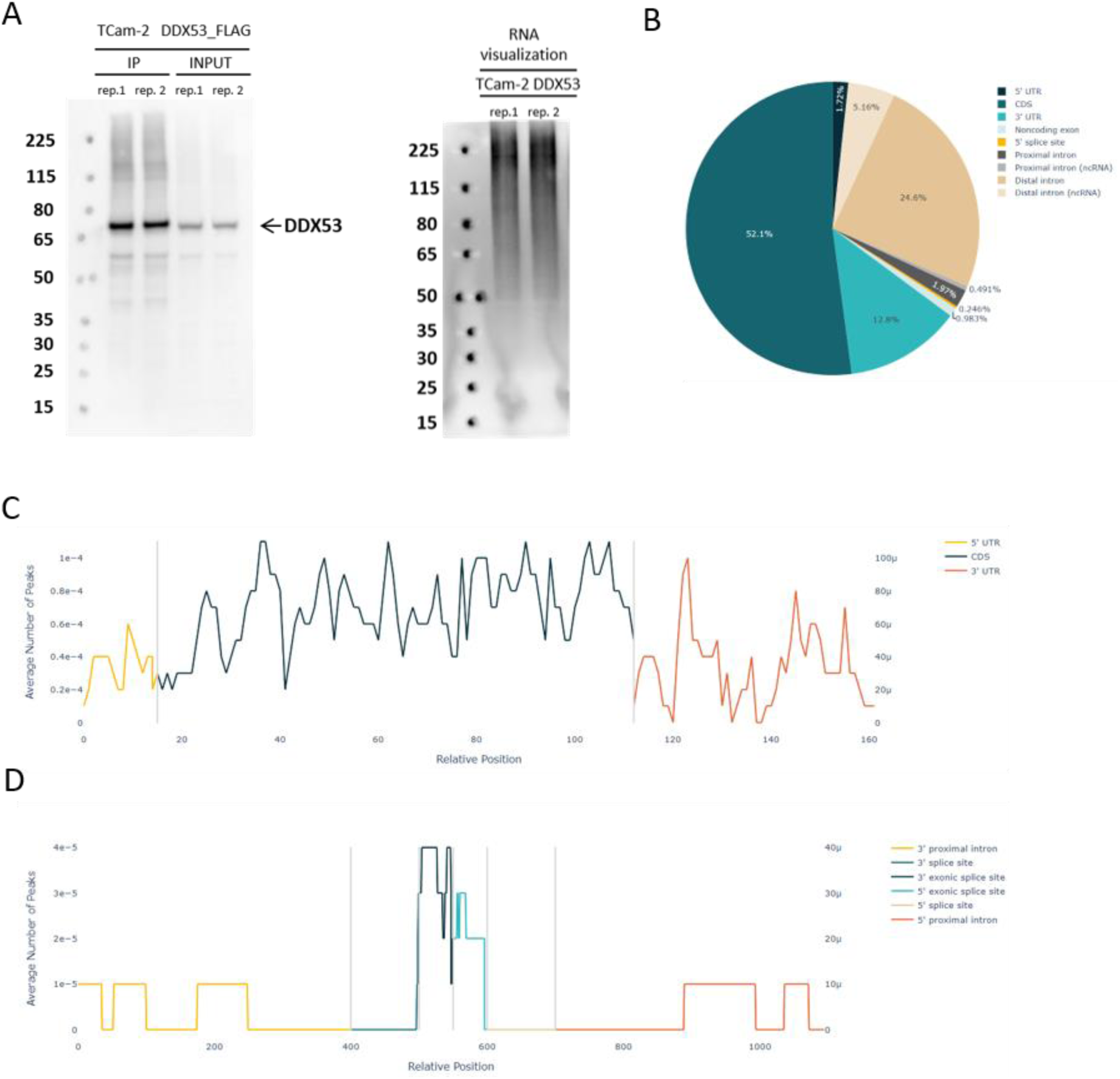
Identification of DDX53-bound RNA transcripts by eCLIP and their global profiling. **(A)** Western Blot analysis using anti-FLAG antibody performed on TCam-2 cells expressing *DDX53-FLAG* (left) and visualization of immunoprecipitated RNA (right) from TCam-2 cells. **(B)** Pie chart showing the relative frequency of peaks that map to each feature type, with a |log2FC| ≥ 3 and adjusted p-value ≤ 0.001. Based on reproducible peaks, identified across DDX53 replicates (Supplementary Table S7). **(C)** Peak metagene plot depicting the average number of peaks mapped to certain genomic regions. The number of peaks was calculated for each region of every gene, the lengths of the regions were normalized, and the average number of peaks for a set number of positions along the regions was calculated. Prepared based on reproducible peaks (|log2FC| ≥ 3 and adjusted p-value ≤ 0.001), identified across DDX53 replicates (Supplementary Table S7). **(D)** Peak metaintron plot depicting the average number of peaks mapped to certain intronic regions. The number of peaks was calculated for each region of every gene, the lengths of the regions were normalized, and the average number of peaks for a set number of positions along the regions was calculated. Prepared based on reproducible peaks (|log2FC| ≥ 3 and adjusted p-value ≤ 0.001), identified across DDX53 replicates (Supplementary Table S7).

**Figure 6.**
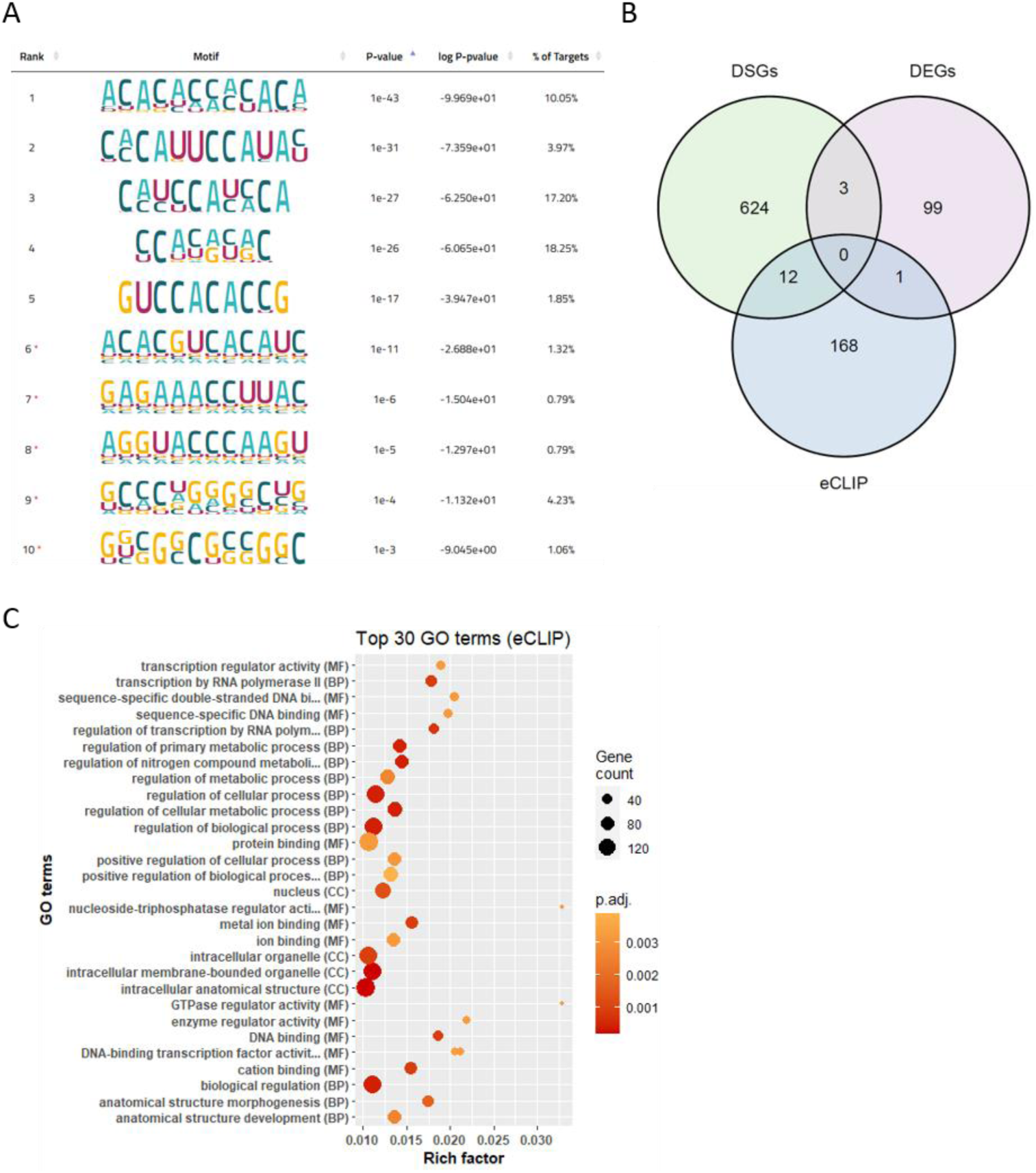
Identification and characterization of DDX53 RNA targets by eCLIP. **(A)** Significant binding motifs of DDX53 binding sites, sorted by p-value and identified by HOMER de novo motif analysis. **(B)** Venn diagram illustrating the total number of DSGs and DEGs based on RNA-seq data in comparison to the list of transcripts detected in DDX53 eCLIP, thus directly bound by DDX53 (Supplementary Table S9). Intersection was prepared using Ensembl IDs. **(C)** Gene ontology (GO) analysis for the transcripts bound by DDX53 based on eCLIP data. The abbreviations in the brackets represent the three GO categories: biological process (BP), molecular function (MF), and cellular component (CC). For the complete list of significant GO terms see Supplementary Table S10 (adjusted p-value < 0.05).

**Figure 7.**
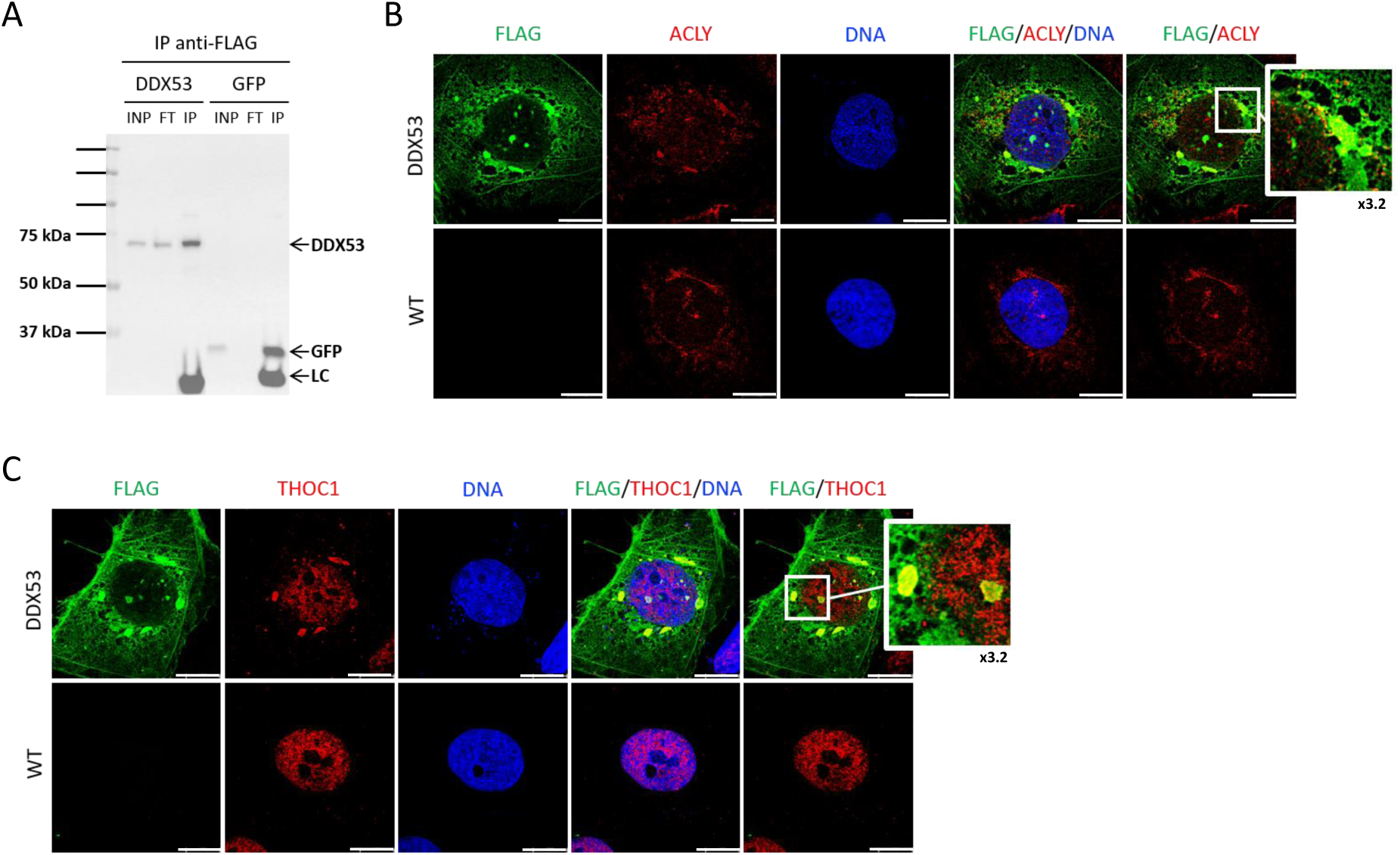
DDX53 potential protein interactors identified by Co-IP-MS. **(A)** Representative Western Blot after immunoprecipitation of FLAG-tagged DDX53 and GFP (as a control) using magnetic beads coated with anti-FLAG M2 antibody. Input (INP, 0.8%), flow-through (FT, 0.8%) and immunoprecipitated (IP, 0.8%) fractions probed with FLAG antibody are shown. **(B, C)** Representative confocal images of TCam-2 cells after transient transfection (DDX53) and WT, which were double immunostained for ACLY and FLAG **(B)** or THOC1 and FLAG **(C)**, respectively. The FLAG antibody detects the DDX53 protein tagged with FLAG sequence. Signal colocalization of DDX53 and its potential protein interactors is shown in the magnified inset (×3.2). Scale bar 15 μm.

### 3.5. DDX53 potential protein interactors are important players during male gamete generation

To identify the protein interaction network of DDX53 we performed co-immunoprecipitation followed by mass spectrometry analysis (Co-IP-MS) on TCam-2 cells with transient overexpression of FLAG-tagged DDX53 and GFP (as control) using FLAG antibody (Figure 7A). This experiment enabled us to identify 35 potential DDX53 interactors (Supplementary Table S11, Supplementary Figure S4A). These proteins represent a diverse group, but members are involved in transcriptional and epigenetic regulation (MTA3, SCML2, SUPT6H), RNA processing and nuclear transport (NUP54, PRCC), and cellular stress response (HSPA9, DNAJB1), collectively contributing to the maintenance of cellular homeostasis. Additional STRING analysis showed that some of the Co-IP-MS–identified proteins are functionally interconnected (Supplementary Table S12). Gene ontology enrichment analysis performed using the whole list of potential DDX53 interactors showed that they are associated with pathways such as citrate metabolic process (ACLY, ACO2) and chaperone cofactor-dependent protein refolding (DNAJB1, HSPA9), processes important in phases of sperm development (Supplementary Figure S4B, Supplementary Table S13) [61,62]. Then, we selected three proteins that are expressed in the human testis at a high level and are present particularly in the germ cells where DDX53 is also expressed (spermatogonia, spermatocytes, and spermatids; according to HPA) and performed further interaction validation using confocal microscopy (Figure 7B-D). Firstly, we took a closer look at the ACLY (also known as ACL/ATPCL), enzyme responsible for the synthesis of acetyl-CoA, important in the context of fatty acids and the acetylation of proteins and histones. Moreover, it has been shown that the ortholog of this gene in Drosophila melanogaster is required for proper male spermatogenesis and that in germ cells of rats ACLY is a potential heat sensitive target [63,64]. The immunofluorescence co-staining showed signal co-localization (Figure 7B), especially at the structures near the nucleus which may indicate that ACLY and DDX53 cooperate during human spermatogenesis. Next, we investigated the interaction of DDX53 with THOC1 (known also as P84/HPR1) and record co-localization signal coming from the structures within and outside the nucleus (Figure 7C). The THOC1 is a subunit of THO complex required for transcription elongation and mRNAs nuclear export [65,66] and has been shown to bind pachytene piRNAs precursors in mouse testis. This feature is especially important in the context of spermatogenesis as pachytene piRNAs emerge when spermatocytes enter the pachytene phase of meiosis [67]. The co-localization of THOC1 and DDX53 underlines the possible role of DDX53 in these processes. It is worth noting that we detected DDX53-positive condensates (Figure 7B,C) and observed colocalization with ACLY and THOC1 in some of these structures. This may suggest that DDX53 and these two interactors participate in germ granules, but this observation requires deeper characterization. Lastly, we investigated the possible interaction of DDX53 with germline-specific Polycomb protein, an epigenetic regulator SCML2 (Supplementary Figure S4C), which has established role in human spermatogenesis supported by other reports [68–70]. Based on co-immunofluorescence staining, we were not able to observe the co-localization of SCML2 and DDX53, thus the potential interaction between these two proteins, based on our Co-IP-MS, requires further validation.

## 4. Discussion

DEAD-box helicases are crucial enzymes involved in various aspects of RNA metabolism. RNA helicases participate in nearly all major RNA-related processes, including RNA modification (such as m⁶A) [71,72] pathways, splicing, RNA degradation, nuclear export, translation, RNP granule dynamics, chromatin–RNA coupling, and RNA quality control [34,73,74]. Their primary function is to remodel structured RNAs and dynamically reorganize RNP complexes, efficient recruitment and activity of RNA-processing enzymes and cofactors. These activities are vital for the proper development of germ cells and sperm maturation [30,73,74]. Although current studies have clarified specific functions of selected DEAD-box helicases in the context of spermatogenesis [75–77], there is a continued need for evaluation of other protein family members to uncover their unknown roles, protein interactors, and targets. Recent reports mentioned the *DDX53* as a new candidate gene possibly associated with spermatogenesis failure based on WGS/WES genetic screening of infertile patients with non-obstructive azoospermia (NOA) [1,2]. The variant identified in infertile male patients results in a missense mutation that alters a residue at the C-terminal end of the KH RNA-binding domain located in the N-terminal region of DDX53. This potentially could modulate RNA substrate specificity as this domain favors single-stranded RNA recognition and binding. Here, for the first time, we describe novel findings on DDX53 in the context of human spermatogenesis, highlighting its role in RNA metabolism. In our research, we used the TCam-2 cells overexpressing *DDX53-FLAG* as an *in vitro* male germline model. This approach enabled us to uncover the coordinated network of RNA targets and protein interactors associated with DDX53 in a cellular environment that mimics early germline status. While the TCam-2 cell line does not entirely capture the full spectrum of germ cell differentiation occurring in the human testis, it remains a useful system for investigating germline-associated mechanisms [37,78]. Therefore, our findings should be regarded as indicative, with emphasis placed on those gene and protein targets that are likewise detected in human male germ cells expressing *DDX53*, as reported by the Human Protein Atlas. Our results demonstrate the importance of DDX53 protein in human transcriptome regulation, which may be compromised in male patients with rare genetic variants of *DDX53* gene.

### 4.1. DDX53 shapes the TCam-2 transcriptome

Firstly, we investigated the influence of DDX53 on TCam-2 cells’ transcriptome by performing RNA sequencing. Although the TCam-2 cell line represents a model of germ cells, it should be noted that these cells originate from a testicular germ cell tumor (seminoma) and harbor multiple cancer-related features. In our study, among the differentially expressed genes (DEGs), we detected the *SERPINF1*, upregulated in TCam-2 cells with DDX53 protein overexpression. SERPINF1 (also known as PEDF) is a multifunctional protein that exerts anti-angiogenic properties, which are especially important given the avascularity of the testicular seminiferous tubules, but also cancer metastasis. In humans, *SERPINF1* is present in testicular tissue, particularly in Sertoli, Leydig, and peritubular cells, and extracellular matrix, but not present in germ cells [28]. Previous studies have suggested its involvement in spermatogenesis [79,80] and underscored its relevance for diverse cancers [81]. Another example of a gene whose expression is positively regulated by DDX53 in TCam-2 cells is the *MZT1* (also known as *MOZART1*), coding a protein that is a component of the gamma-tubulin large complex, primarily localized to the centrosome and spindle. Although *MZT1* has not yet been studied in human spermatogenesis, it was reported that *Drosophila* Mzt1 is required to maintain sperm motility and male fertility during aging [82]. In theory, DDX53 may underlie proper *MZT1* regulation, functionally significant also for human male germ cells differentiation.

### 4.2. DDX53 modulates alternative splicing of germline-relevant genes

In this work, we identified multiple differentially spliced genes (DSGs) upon *DDX53* overexpression. Splicing alterations were more prominent than changes in overall gene expression, which aligns with the anticipated role of DDX53 in RNA metabolism, rather than transcriptional regulation. Among the DSGs was the *LRRC8A*, coding for a component of the volume-regulated anion channel, which helps to maintain cell volume homeostasis and has fundamental significance for the physiological function of sperm. Knockout of *Lrrc8a* in mice led to defective development of spermatids and male infertility [83]. Our eCLIP results indicate that DDX53 binds *LRRC8A* mRNA within CDS and 3’UTR, raising the possibility that sequence variants in DDX53 may perturb *LRRC8A* mRNA processing, with potential consequences for protein function and male germ cell biology. Another interesting transcript, bound by DDX53 within 3’UTR, and classified as differentially spliced in TCam-2 cells upon *DDX53* overexpression, is *TFAM*. TFAM is a key mitochondrial transcription factor, essential not only for the transcription but also maintenance of mitochondrial DNA (mtDNA). This protein reaches the highest expression level in spermatids (similarly to DDX53), and dysregulation of *TFAM* expression has been correlated with qualitative impairment of spermatogenesis [84]. Studies have shown that downregulation of TFAM protein levels leads to a decrease in mtDNA copy number during spermatogenesis, which is believed to be a mechanism preventing the transmission of paternal mtDNA to the offspring, ensuring predominant maternal inheritance of mitochondria [85,86].

The overlap between the number of potential DDX53-induced DSGs and identified DDX53-bound transcripts was relatively low (according to RNA-seq and eCLIP data comparison), but it is important to remember that DDX53 may influence alternative splicing of many key genes, important for spermatogenesis, without direct binding to their mRNA. It is well-known that proteins may influence AS by interacting with other regulatory proteins, rather than by directly binding to RNA transcripts. Notably, multiple proteins involved in alternative splicing have been found among the potential DDX53 protein interactors as shown by Co-IP-MS, such as CELF2 [87], RBM26 [88], and THOC1 [89]. We performed immunofluorescence co-staining for DDX53 and THOC1 and observed signal overlap, which indicates a possible functional interplay. Another noteworthy example of a gene classified as differentially spliced upon *DDX53* overexpression, but not directly bound by DDX53, is the *SRSF1*. SRSF1 protein is expressed in various stages of male germ cell development. A recent study showed that it is essential for the proper expression and alternative splicing of spermatogonia-related genes in mouse testis [90]. Moreover, conditional knockout (cKO) mice with SRSF1 deficiency were infertile and exhibited a Sertoli cell-only syndrome (SCOS) phenotype. Another group showed that SRSF1 loss led to meiotic arrest at the pachytene stage, emphasizing that germ cell-specific *Srsf1* knockout results in spermatogenesis failure and thus male infertility [91]. Interestingly, the SEPTIN7 also emerged among the DSGs. This gene codes for a cytoskeletal GTPase that plays a crucial role in maintaining cell shape and structural integrity during spermatogenesis. SEPTIN7 is robustly expressed throughout all stages of human spermatogenesis. This protein migrates from the midpiece to the annulus as spermatozoa mature, suggesting that it may be involved in regulating sperm maturation [92]. The annulus is a septin-based ring structure at the junction of the sperm midpiece and principal piece that functions as a diffusion barrier and is required for proper flagellar organization and sperm motility. Proper regulation of *SEPTIN7* expression may be critical for preventing sperm defects, highlighting a potentially important, although secondary, role of DDX53.

Spermatogenesis relies on highly specialized RNA processing programs, reflected by the exceptionally high prevalence of alternative splicing observed in the testis, a feature shared with the human brain [93,94]. In this context, recent studies have demonstrated DDX53 expression in both tissues and its role in neuronal function [3,4], suggesting that DDX53 may contribute also to complex RNA regulatory mechanisms in male germ cells. Despite the widespread occurrence of alternative splicing in the testis, most transcript isoforms remain poorly annotated and functionally uncharacterized [95], underscoring the need to define RNA-binding regulators such as DDX53.

### 4.3. DDX53 binds numerous germline-associated transcripts

Our findings support DDX53 as an upstream regulator of RNA metabolism based on its ability to bind a broad range of RNA molecules in human TCam-2 cells, as demonstrated by eCLIP-seq data including transcripts already discussed in the context of differentially spliced genes (DSG). Among the RNA targets of the DDX53, numerous transcripts coding proteins and previously recognized as important in the context of spermatogenesis were identified. Below, we discuss selected examples highlighting the potential role of DDX53 in male germ cells. An important example of mRNA targeted by DDX53, relevant in the context of spermatogenesis, is the *TAF7* (bound within its coding sequence). This gene codes for spermatogenic transcription regulator whose mutations increase the risk of azoospermia [96]. The intracellular localization of TAF7 is dynamically regulated, as it is present in the nucleus in developing germ cells until the late pachytene stage, after which it becomes cytoplasmic [97]. Precise regulation of *TAF7* mRNA may be a key to its proper testis-specific expression pattern, which could be disrupted in patients with *DDX53* mutations, leading to infertility. Another noteworthy example is the LIN28B, an RNA-binding protein that acts as a post-transcriptional regulator of genes involved in developmental timing and self-renewal through its interaction with miRNAs [98]. In humans, LIN28B is mainly expressed in the testis and placenta. Moreover, within the adult human testis, *LIN28B* transcripts are found in the germ cells and not in the somatic cells. A recent study suggested that LIN28B may act as a proliferation driver in the most undifferentiated spermatogonia, and that a decrease in its expression could impair spermatogenic capacity, resulting in oligozoospermia in adult men [98]. DDX53 binding (within 3’UTR) to *LIN28B* mRNA suggest that DDX53 may act upstream of LIN28B-mediated pathways. Of particular interest, *NCBP2* was identified as a potential mRNA target of DDX53. The product of this gene plays a vital role in mRNA processing events through its binding to the 5’ cap of nascent pre-mRNAs. Results from single-cell RNA sequencing of testicular cell samples from NOA patients indicated that NCBP2 is positively associated with spermatogenesis [99]. Although its precise role during male germ cell development has not yet been investigated, the NCBP2 has been determined as a significant predictor of non-obstructive azoospermia (NOA) [99]. An important example might be also the *BRD3* gene, whose mRNA is bound by DDX53 within 3’UTR. In mice, *Brd3* is enriched in round spermatids, demonstrating a tightly regulated expression pattern in the male germ cell lineage and suggesting its specific role for spermatogenesis [100]. While the precise role of BRD3 in spermatogenesis is yet unknown, its membership in the bromodomain and extraterminal (BET) family proteins implies involvement in transcriptional regulation during germ cell development. BET proteins, including BRD3, are known to interact with acetylated histones, influencing chromatin structure and gene expression. The specific enrichment of *BRD3* transcripts in round spermatids points to a potential role in the transcriptional programs necessary for the maturation of spermatids into spermatozoa. In humans, *BRD3* transcripts are expressed in the spermatogonia, spermatocytes, and spermatids, but the protein was specifically observed in spermatids [101]. Additionally, DDX53 protein also binds the mRNAs of *LIN7C* and *DLC1*, both within coding sequence. These proteins exhibit stage-specific expression in germ cells during spermatogenesis, which implies their testis-specific functions. Lin7c has been proposed as a potential marker of early spermatogenesis [102], and a role for DLC1 in chromatin condensation and spermatid shaping has been suggested [103]. Although more studies are needed to elucidate their precise function in the testis, it has been suggested that they are important in germ cell development. Based on eCLIP data, it is possible that DDX53 governs the stage-specific expression of these genes in the human testis. Importantly, eCLIP analysis also revealed that DDX53 binds to its own mRNA. Although this finding does not directly support a self-regulatory effect, it is consistent with feature commonly observed among RNA-binding proteins [58–60] and may suggest a potential autoregulatory feedback mechanism, which remains to be functionally validated.

### 4.4. Protein interactome supports an RNA-centric regulatory role

Analysis of the STRING interaction network based on DDX53 potential protein interactors identified by Co-IP-MS) suggests that DDX53 operates within an integrated transcriptional and post-transcriptional regulatory framework. The interactome comprising POLR2B, SUPT6H, RPRD1A, THOC1, NUP54 and NUP210 forms a coherent axis spanning transcription by RNA polymerase II, co-transcriptional RNA maturation, and nuclear export of messenger ribonucleoprotein complexes. Among the potential DDX53-protein interactors, there are proteins involved in pre-mRNA splicing and RNP dynamics. Collectively, our data suggest that DDX53 is involved in mediating mRNA metabolism, which is critical for precisely controlled germline-specific gene expression.

### 4.5. DDX53-positive condensates suggest potential involvement in germline-specific granules

We investigated DDX53 subcellular localization, demonstrating its presence mostly in the cytoplasm with partial nuclear localization in TCam-2 cells. Although several studies have detected positive DDX53 ChIP-based signals [15,16,23,27,54] and we confirmed the partial nuclear localization of DDX53 in TCam-2 cells, our ChIP-seq analysis did not support direct DNA binding by DDX53 (unpublished data). Based on this, previously reported positive ChIP assay results for DDX53 should be interpreted with caution. While nonspecific antibody interactions may partially account for these observations, our data do not preclude an indirect or transient association of DDX53 with chromatin, potentially mediated through protein–protein interactions.

To further clarify the nuclear context of DDX53, we examined its intracellular organization and observed distinct DDX53-positive condensates in TCam-2 cells that resemble germline-type RNP granules. These specialized ribonucleoprotein (RNP) granules are a hallmark of all germ cells and are composed of germline-specific proteins and RNAs [104,105]. Although over recent decades various types of RNA granules have been identified in germ cells across species, still very little is known about these germline-specific condensates in humans. In the *Drosophila* embryo, nuclear germ granules were shown to promote germ cell divisions thereby increasing PGC number for the next generation [106]. Furthermore, one of the conserved components of germ granule proteins is Ddx4 (known also as Vasa) [56,106]. Although not expressed in TCam-2 cells [107], DDX4 is an evolutionarily conserved germline protein that belongs to the DEAD-box family of RNA helicases, as does DDX53. Importantly, the identification of DDX53-positive condensates remains preliminary and requires further validation using canonical germ granule markers and additional experimental approaches, particularly under physiological conditions. This may represent a valuable direction for future research and could provide additional insight into the role of DDX53 in the human germline.

## 5. Conclusions

Collectively, the presence of DDX53 in human testicular tissue, together with its influence on transcriptome in TCam-2 cells and its capacity to bind a broad repertoire of RNAs, implies its role in male germ cell biology. Moreover, our data suggest that DDX53 protein may affect various testis-specific post-transcriptional gene regulation mechanisms that influence male germ cell differentiation. Further research is warranted to elucidate the precise mechanisms by which DDX53 influences spermatogenesis in diverse stages of germ cell development, but this study highlights the relevance of DDX53 as an upstream regulator of RNA metabolism, vital for proper germline development.

## Author Contributions

Conceptualization: N.R., A.M.; Methodology: A.J.B., K.T., A.M., T.K.; Investigation: A.J.B., K.T., A.M., M.D.; Formal analysis: E.D., A.J.B., J.Z.-W. (NGS data); E.I., D.C. (MS data); T.K.; Resources: M.K., M.O., A.Y. (selection of gene of interest based on previous studies), M.H.; Data curation: M.K., M.O., A.Y., M.H.; Writing – original draft: A.J.B.; Writing – review & editing: A.J.B., N.R., K.T., E.I., A.M., T.K., M.D., E.D., J.Z.-W., M.O., D.C., M.H., A.Y., M.K.; Supervision: N.R.; Funding acquisition: N.R. All authors have read and agreed to the published version of the manuscript.

## Institutional Review Board Statement

Not applicable.

## Informed Consent Statement

Not applicable.

## Data Availability Statement

Sequencing data have been deposited in Gene Expression Omnibus (GEO, https://www.ncbi.nlm.nih.gov/geo/) under accession code GSE312193 (RNA-seq) and GSE312179 (eCLIP-seq). The datasets generated in this study are available in Zenodo: https://doi.org/10.5281/zenodo.17815115.

## Supporting information

Supplementary Material DDX53

## Acknowledgments

The scientific work was created with the assistance of the infrastructure of Department of Cellular, Computational and Integrative Biology (CIBIO, University of Trento) and Poznan Supercomputing and Networking Center (PSNC). We are grateful to Augustyn Moliński for supporting the investigation of DDX53 localization and for expert assistance with confocal microscopy. We thank Marcin Sajek for help with NGS data deposition and Katarzyna Kiwerska for help with IHC experiment. This manuscript has been released as a preprint at bioRxiv (https://doi.org/10.64898/2026.02.03.703580), available under a CC-BY-NC-ND license.

## Abbreviations

RNA-seq: RNA sequencing
eCLIP: Enhanced crosslinking and immunoprecipitation
Co-IP-MS: Co-immunoprecipitation coupled with Mass Spectrometry
WES: Whole-exome sequencing
WGS: Whole-Genome sequencing
CTAs: Cancer-testis antigens
NOA: Non-obstructive azoospermia
HPA: Human Protein Atlas
hPGCs: Human primordial germ cells
ChIP: Chromatin Immunoprecipitation
IHC: Immunohistochemistry
DEGs: Differentially expressed genes
AS: Alternative splicing
DSGs: Differentially spliced genes
GO: Gene ontology
SCOS: Sertoli cell-only syndrome

## Supplemental Tables

Supplementary Table S1 Normalized counts (CPM) based on RNA-seq of TCam-2 cells.

Supplementary Table S2 Differentially expressed genes (DEGs) based on RNA-seq of TCam-2 cells.

Supplementary Table S3 Normalized transcript-level values (TPM) for TCam-2 cells quantified with Salmon from RNA-seq data.

Supplementary Table S4 Differentially spliced genes (DSGs) based on SUPPA2 analysis of RNA-seq data.

Supplementary Table S5 Genes overlap results for DSGs, DEGs, and genes involved in male gamete generation.

Supplementary Table S6 Gene Ontology and pathway enrichment analysis of DSGs detected upon DDX53 over-expression.

Supplementary Table S7 Detailed information on detected DDX53 eCLIP peaks.

Supplementary Table S8 List of transcripts bound by DDX53 identified by eCLIP.

Supplementary Table S9 Genes overlap results for DSGs, DEGs, and DDX53 eCLIP targets.

Supplementary Table S10 Gene Ontology analysis of DDX53 eCLIP targets.

Supplementary Table S11 Results of DDX53 co-immunoprecipitation followed by mass spectrometry.

Supplementary Table S12 DDX53 protein-protein interaction network analysis performed with the STRING database.

Supplementary Table S13 Gene Ontology enrichment analysis based on proteins identified in DDX53 co-immuno-precipitation experiment.

Supplementary Table S14 List of primers.

