## Supplementary Material DDX53 for "The RNA helicase DDX53 (CAGE) contributes to RNA metabolism in a human germ cell model"

[1] Institute of Human Genetics, Polish Academy of Sciences, Poznan, Poland
[2] Department of Gene Expression, Institute of Molecular Biology and Biotechnology, Adam Mickiewicz University, Poznan, Poland
[3] Department of Mathematical and Statistical Methods, Poznan University of Life Sciences, Poznan, Poland
[4] Department of Cellular, Computational and Integrative Biology (CIBIO), University of Trento, Trento, Italy
[5] DNRF Center for Chromosome Stability, University of Copenhagen, DK-2200 Copenhagen, Denmark
[6] Laboratory of Mass Spectrometry, Institute of Biochemistry and Biophysics, Polish Academy of Sciences, Warsaw, Poland
[7] Department of Urology, Poznan University of Medical Sciences, Poznan, Poland
[8] Department of Obstetrics, Gynecology, and Reproductive Sciences, Magee-Womens Research Institute, University of Pittsburgh School of Medicine, Pittsburgh, PA, United States
[9] Department of Pathology, University of Pittsburgh School of Medicine, Pittsburgh, PA, United States
[10] Department of Genetics, University of Pittsburgh School of Public Health, Pittsburgh, PA, United States

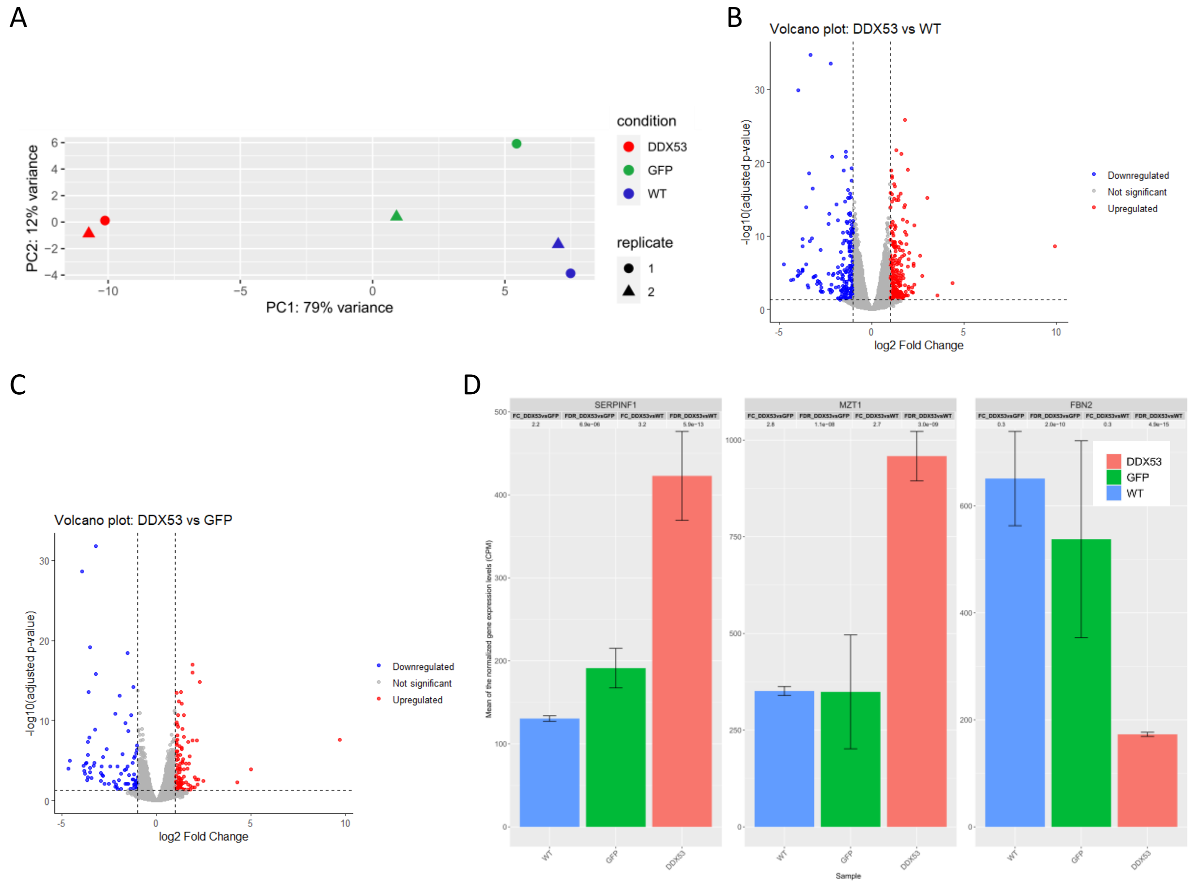


Supplementary Figure S1. Principal component analysis results of TCam-2 RNA-seq data, volcano plots, and expression results for three DEGs (SERPINF1, MZT1, and FBN2) selected for validation via qRT-PCR. (A) Principal component analysis (PCA) of TCam-2 RNA-seq samples (Supplementary Table S1). The TCam-2 with DDX53 overexpression samples are highlighted in red and the GFP and WT are highlighted in green and blue, respectively. Test samples (DDX53) separate from the control (GFP and WT) samples along principal component 1 (PC1). The two biological replicates are presented with different shapes. (B) Volcano plot of DEGs identified using DESeq2 from RNA-seq data of DDX53-overexpressing TCam-2 cells versus WT (Supplementary Table S2). Dashed lines represent twofold change in expression (vertical lines) and adjusted FDR < 0.05 cutoff (horizontal line). (C) Volcano plot of DEGs identified using DESeq2 from RNA-seq data of DDX53-overexpressing TCam-2 cells versus GFP (Supplementary Table S2). Dashed lines represent twofold change in expression (vertical lines) and adjusted FDR < 0.05 cutoff (horizontal line). (D) Barplots showing the mean expression level (CPM) of SERPINF1, MZT1, and FBN2 genes in TCam-2 cells with stable expression of DDX53 (red) and GFP (green) as well as WT (blue) based on RNA-seq analysis (Supplementary Table S1, 2). Additionally, fold change (FC) and adjusted p-value (FDR) for both comparisons (DDX53 vs. GFP and DDX53 vs. WT) are presented in the upper part of the plots. Error bars in black represent the standard deviation (SD) calculated for the CPM expression values from the two biological replicates.

A

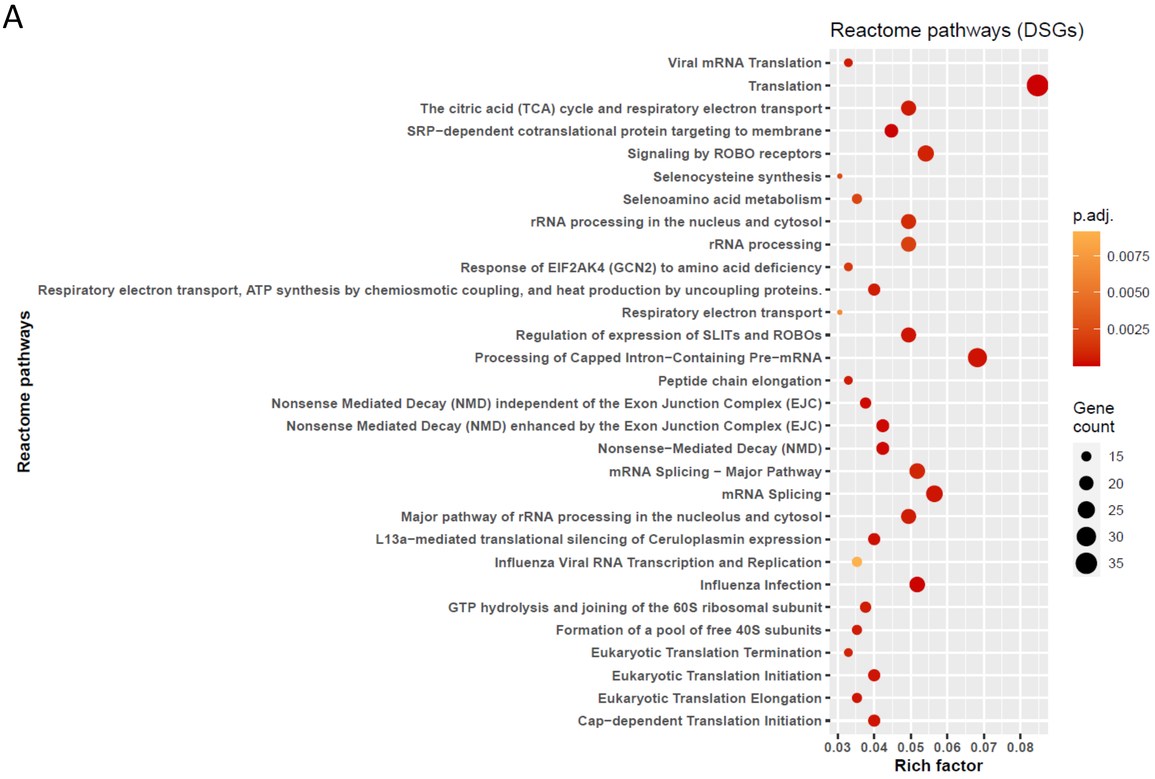


Supplementary Figure S2. Potential pathways regulated by DSGs detected upon DDX53 overexpression and RNA-seq. (A) Reactome analysis for the DSGs based on the RNA-seq data. For the complete list of significant Reactome terms see Supplementary Table S6 (adjusted p-value < 0.05).

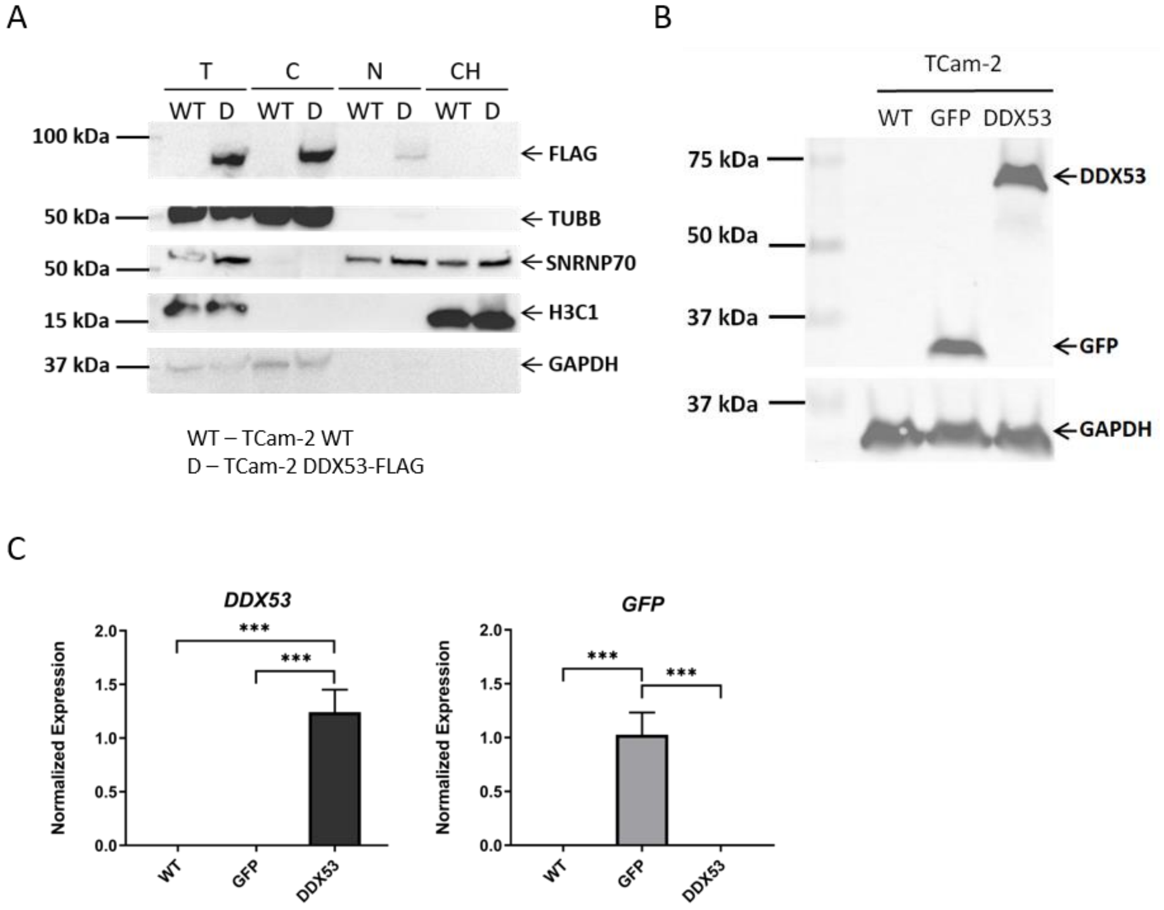


Supplementary Figure S3. Subcellular fractionation and validation of FLAG-tagged DDX53 and GFP expression in TCam-2 cells following transient transfection. (A) Western blot of TUBB (cytoplasmic marker), SNRNP70 (nuclear marker), and H3C1 (chromatin marker) as well as FLAG; following cellular fractionation of TCam-2 cells with DDX53-FLAG overexpression (denoted as D) and WT cells. The appropriate subcellular fractionation was confirmed by enrichment of the respective marker in each fraction. (B) Representative Western Blot analysis for proteins tagged with FLAG (DDX53 and GFP) on TCam-2 cells after transient transfection and wild type (WT, as control) using FLAG antibody. GAPDH was used as a loading control. (C) The mRNA expression level of the DDX53 and GFP with GAPDH and ACTB as reference genes. The TCam-2 cells were assayed 48h after transfection. Error bars represent standard deviation (SD) with n = 3, ***$p < 0.001$. The p values were calculated by ordinary one-way ANOVA non-parametric test with Tukey's multiple comparisons test.

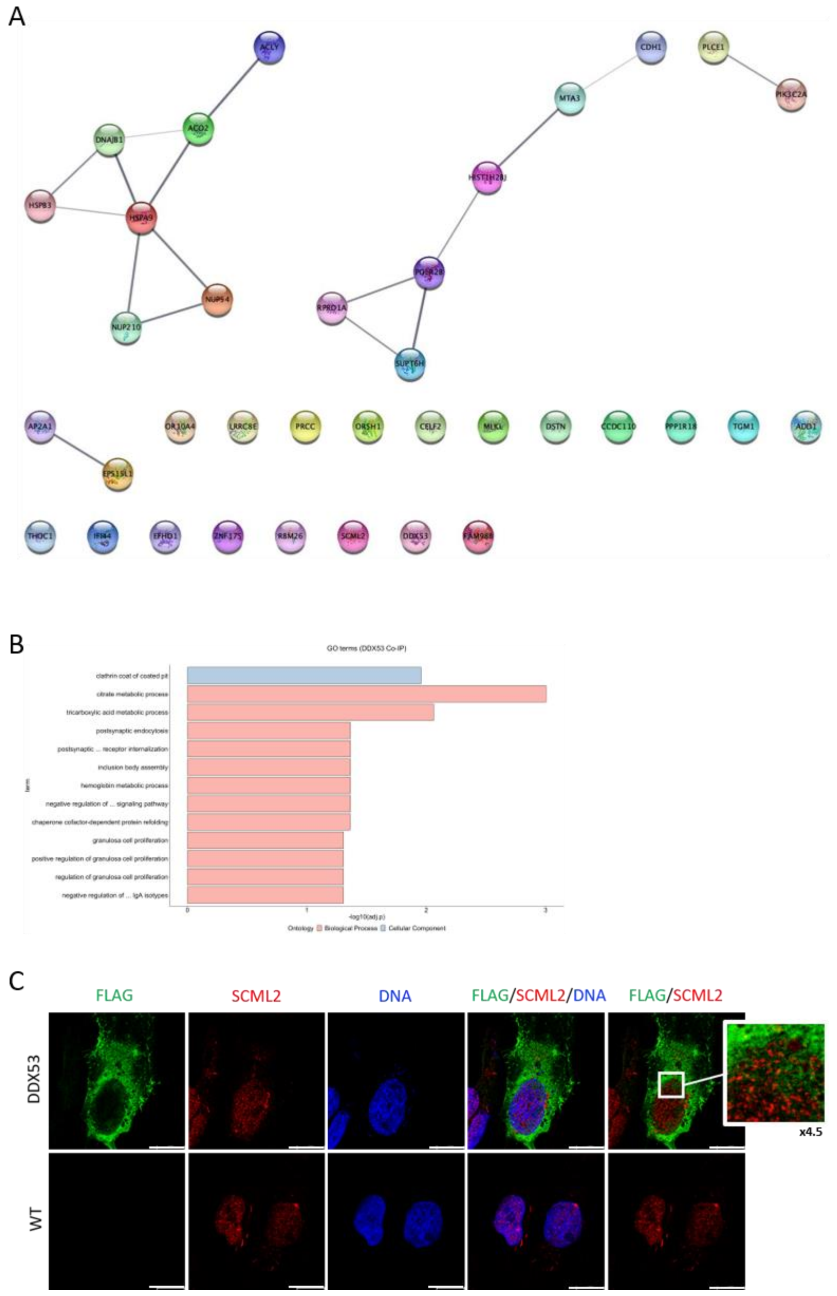


Supplementary Figure S4. Identification of potential protein interactors for DDX53 using Co-IP-MS. (A) Protein-protein interaction (PPI) network created among 35 identified potential protein interactors and DDX53, performed using the STRING database (Supplementary Table S12). The 0.4 cutoff score was used to filter out interactions below this value and the thickness of grey edges between proteins indicates how high was the score of interaction. (B) Gene ontology (GO) analysis of potential protein interactors of DDX53 identified by Co-IP-MS (adjusted p-value $\leq 0.05$, Supplementary Table S13). The color represents two GO categories (blue: CC, red: BP). (C) Representative confocal images of TCam-2 cells after transient transfection (DDX53) and WT, which were double immunostained for SCML2 and FLAG. The FLAG antibody detects the DDX53 protein tagged with FLAG sequence. Signal colocalization for DDX53 and SCML2 was not observed, as presented in the magnified inset (×4.5). Scale bar 15 µm.
